# Cargo selective vesicle tethering: the structural basis for binding of specific cargo proteins by the Golgi tether component TBC1D23

**DOI:** 10.1101/2023.11.22.568253

**Authors:** Jérôme Cattin-Ortolá, Jonathan G. G. Kaufman, Alison K. Gillingham, Jane L. Wagstaff, Sew-Yeu Peak-Chew, Tim J. Stevens, David J. Owen, Sean Munro

**Affiliations:** MRC Laboratory of Molecular Biology, Francis Crick Avenue, Cambridge CB2 0QH, UK; Cambridge Institute for Medical Research, Cambridge Biomedical Campus, Hills Road, Cambridge CB2 0XY, UK

## Abstract

For accurate membrane traffic it is essential that vesicles and other carriers tether and fuse to only the correct compartment. The TGN-localised golgins golgin-97 and golgin-245 capture transport vesicles arriving from endosomes via the protein TBC1D23 that forms a bridge between the golgins and endosome-derived vesicles. The C-terminal domain of TBC1D23 is responsible for vesicle capture, but how it recognises a specific type of vesicle was unclear. A search for binding partners of the C-terminal domain surprisingly revealed direct binding to carboxypeptidase D (CPD) and syntaxin-16, both known cargo proteins of the captured vesicles. Binding is via a TLY-containing sequence present in both proteins. A crystal structure reveals how this “acidic TLY motif” binds to the C-terminal domain of TBC1D23. An acidic TLY motif is also present in the tails of other endosome-to-Golgi cargo, and these also bind TBC1D23. Structure-guided mutations in the C-terminal domain that disrupt motif binding in vitro also block vesicle capture in vivo. Thus, TBC1D23 attached to golgin-97 and golgin-245 captures vesicles by a previously undescribed mechanism: the recognition of a motif shared by cargo proteins carried by the vesicle.

**One sentence summary:** A class of transport vesicle destined for the Golgi is recognized by a tether binding directly to the cargo it is carrying.

## Introduction

The transport of proteins between the organelles of the secretory and endocytic pathways is mediated by tubular/vesicular carriers that bud off donor organelles and then fuse with destination organelles. The process by which the vesicles recognise their correct destination involves tethering factors that mediate the initial capture the vesicle prior to the SNARE proteins in the vesicle and destination membranes forming a complex that then drives membrane fusion (*1*, *2*). Depending on the transport step, these tethering factors are long coiled-coil proteins or large protein complexes that are located to a specific organelle (*3–5*). In the case of the Golgi apparatus these tethers include a set of long coiled-coil proteins called ‘golgins’. They are located to specific parts of the Golgi stack via C-terminal domains that either bind small GTPases or form a single transmembrane domain that spans the Golgi bilayer (*6–8*). Their role as tethers has been clearly demonstrated by the finding that replacing their C-terminal domains with one that directs targeting to mitochondria results in the accumulation of specific Golgi-destined carriers at this ectopic location (*9–11*).

The ability of golgins to capture vesicles raises the question of how these and other tethers can recognise the correct vesicles. In the case of the golgins, conserved N-terminal regions are necessary and sufficient to mediate vesicle capture (*10*). Three golgins, golgin-97, golgin-245 and GCC88 can capture endosome-to-Golgi carriers, and the first two share a closely related vesicle capture motif at the N-terminus. This motif binds to a cytosolic protein, TBC1D23, a member of the TBC family of Rab GAPs that is unlikely to have GAP activity as it lacks the conserved catalytic residues necessary for stimulating GTP hydrolysis (Fig. 1A) (*12*, *13*). TBC1D23 is located to the Golgi in a manner that is dependent on the presence of golgin-97 and golgin-245, and it can by itself capture vesicles when relocated to mitochondria indicating that it forms a bridge between the golgins and endosome-derived vesicles (*12*). In addition to the TBC domain, TBC1D23 has a rhodanese domain, and a C-terminal domain that is related to PH domains (*13*)(*14*). The TBC/rhodanese N-terminal region of the protein binds directly to the N-terminus of golgin-97 or golgin-245, and the C-terminal domain is necessary and sufficient for vesicle capture (*12*, *14*). Between these two regions there is a linker region, part of which binds to a complex of three proteins FAM91A1, WDR11, and C17orf75 (*12*, *15*, *16*). The function of this complex is unknown, and the part of TBC1D23 that it binds is not required for vesicle capture (*12*).

**Fig. 1.**
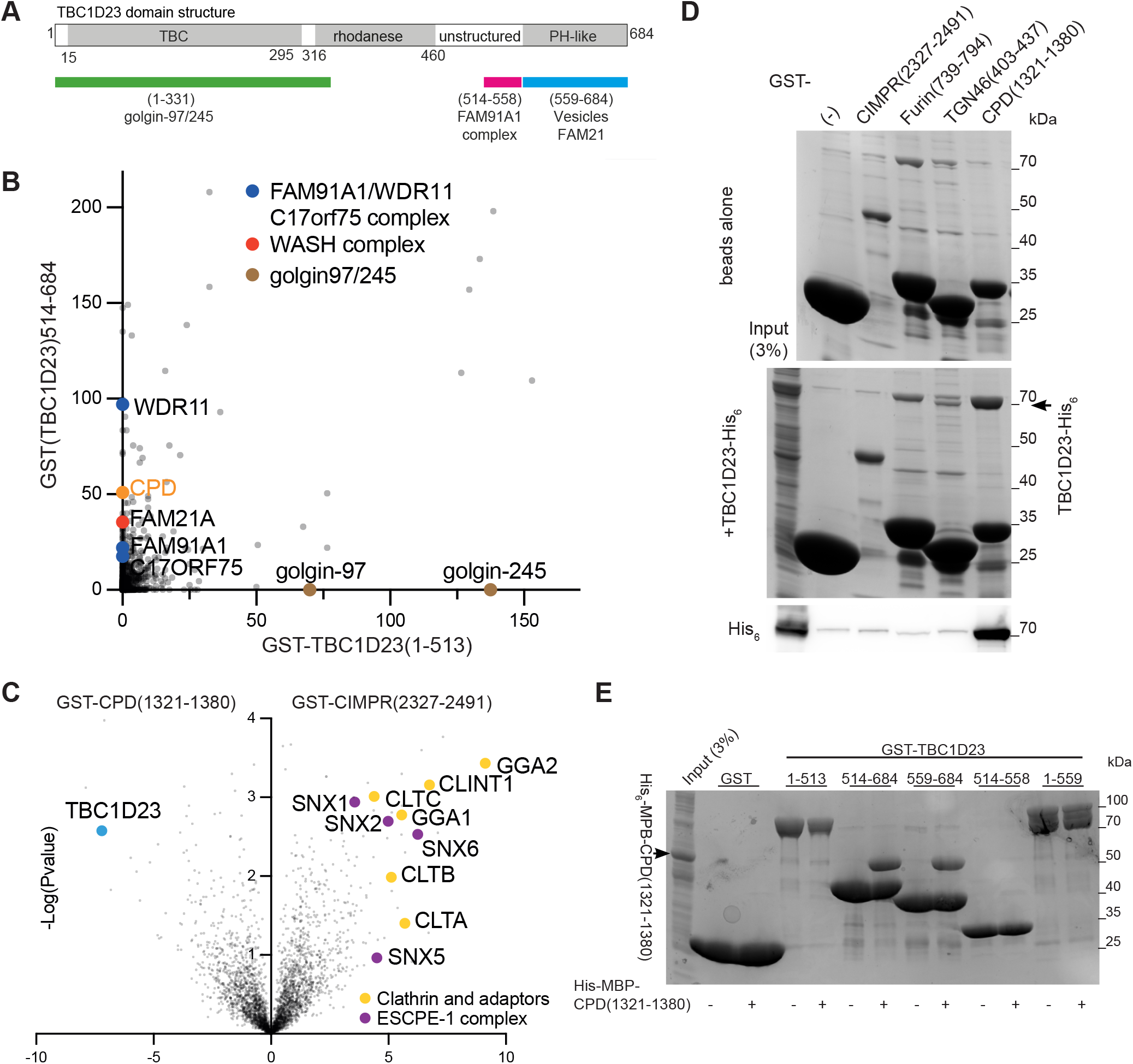
The C-terminal domain of TBC1D23 forms a complex with the cytoplasmic tail of carboxypeptidase D (CPD). (A) Domain structure of mouse TBC1D23 (UniProt Q8K0F1), the version of the protein used in this study. Labelling indicates the regions found by deletion mapping to bind the golgins, the FAM91A1 complex, the WASH complex and vesicles (*12*, *14*). (B) Mass spectrometry analysis from affinity chromatography of 293T cell lysates using GST-TBC1D23 fragments. The plot compares the average spectral counts from two independent replicates of GST-TBC1D23(1-513) versus GST-TBC1D23(514-684). Values are in data S1. (C) Volcano plot of mass spectrometry analysis comparing the eluates from affinity chromatography of 293T cell lysates using cytoplasmic tails of CPD and CIMPR. The plot compares the spectral intensities from proteins bound to each bait, using data from three independent biological replicates. Values are in data S1. (D) Coomassie-stained gel and immunoblot showing that TBC1D23-His6 binds directly and specifically to the cytoplasmic domain of CPD. GST-tagged cytoplasmic domains of several endocytic cargoes were immobilised on beads and incubated with bacterial lysate containing TBC1D23-His_6_. (E) Coomassie-stained gel showing that the C-terminal domain of TBC1D23 is necessary and sufficient for binding to CPD. GST tagged fragments of TBC1D23 were immobilised on beads and incubated with a bacterial lysate containing the cytoplasmic domain of CPD (His6-MBP-CPD(1321-1380)).

The ability of the C-terminal domain of TBC1D23 to capture a specific class of vesicles means that it must recognise a feature shared by these vesicles. Previous studies have shown that the C-terminus can bind to the endosomal WASH complex via the FAM21 subunit which has a long C-terminal tail that also binds several other WASH interactors and regulators (*12*, *17*, *18*). However, WASH is thought to be involved in retromer and retriever-based carrier formation at endosomes, including carriers that are destined for the Golgi and others destined for the plasma membrane (*19*, *20*). This led to the suggestion that TBC1D23 is able to bind to an additional factor that allows it to specifically capture carriers destined for the Golgi (*12*). In this paper we describe the application of an unbiased affinity chromatography approach to identify interaction partners of the C-terminal domain of TBC1D23 and the report that it can bind directly to the cytoplasmic tails of several of cargo proteins that are found in endosome-to-Golgi carriers. This provides a mechanism by which TBC1D23 and its associated golgins can capture and tether endosome-derived carriers as they arrive at the Golgi.

## Results

### TBC1D23 binds directly to the cytoplasmic tail of the endocytic cargo carboxypeptidase D (CPD)

To identify binding partners for the C-terminal PH-like domain of TBC1D23, we immobilised on beads a GST fusion to the domain and performed affinity chromatography of lysates from 293T cells. Mass-spectrometry of the bound proteins revealed, as expected, that the C-terminal domain could selectively enrich the subunits of the FAM91A1 complex along with the WASH complex subunit FAM21, whilst the N-terminal domain bound to its partner golgins (Fig. 1B). As is typical for GST fusion-based purifications, the proteins bound to the C-terminal domain contained several abundant chaperones and cytosolic enzymes that are likely to reflect non-specific interactions, but in addition to these we identified carboxypeptidase D (CPD). This metalloprotease is broadly expressed and removes C-terminal basic residues following the action of furin and related proteases on diverse substrates including neuropeptides and growth factors (*21*, *22*). CPD is known to be primarily localised to the TGN and to recycle through endosomes and the plasma membrane. CPD is a type I protein with a 60-residue cytoplasmic tail, and so to validate this interaction we used a GST fusion to the tail for affinity chromatography from 293T cell lysates. This showed that TBC1D23 has a strong preference for the tail of CPD over that of a control protein - the cation-independent mannose 6-phosphate receptor (CI-MPR), another abundant protein that recycles between endosomes and the TGN (Fig. 1C). The interaction between TBC1D23 and CPD could be recapitulated with proteins expressed in *E. coli*, demonstrating that it is direct (Fig. 1D), and use of truncated forms confirmed that it is the C-terminal domain of TBC1D23 that binds to the CPD tail (Fig. 1E). Taken together, these results show that the cytoplasmic tail of CPD can form a stable complex with residues 559-684 of TBC1D23, the C-terminal domain that mediates vesicle capture.

### TBC1D23 is required for normal trafficking of CPD

To test whether TBC1D23 is required for the trafficking of CPD we used CRISPR-Cas9 to remove the TBC1D23 gene from both human 293 cells and the rat insulinoma line INS-1 (Fig. 2A and fig. S1A). CPD cycles between endosomes and the Golgi, and for some such proteins it has been found that perturbation of retrieval from endosomes results in their destabilisation (*12*, *21*, *23*). In both knockout cell lines, the steady-state level of CPD was reduced, and this could be rescued by expression of TBC1D23-GFP (Fig. 2, A and B). In rat INS-1 cells, it was possible to detect CPD by immunofluorescence, and the levels of the protein in the TGN were reduced in the absence of TBC1D23 (Fig. 2C). Again, this phenotype was rescued by expression of TBC1D23-GFP but not a form lacking the C-terminal domain (Fig. 2, D and E, and fig. S1, B to D).

**Fig. 2.**
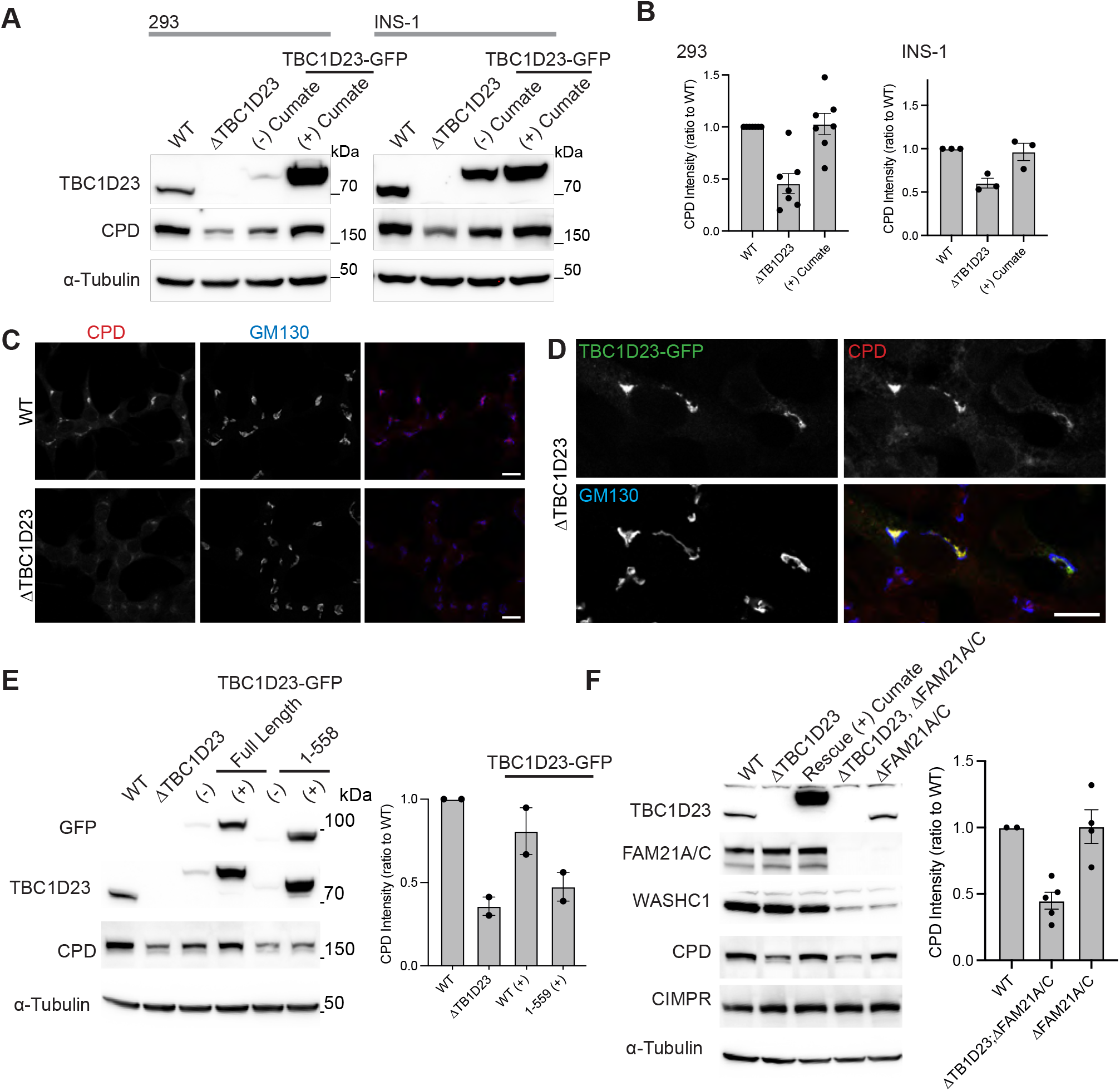
Normal CPD trafficking requires the C-terminal domain of TBC1D23 but does not require the WASH complex. (A) Immunoblots comparing TBC1D23, CPD and α-tubulin in whole cell lysates of 293 and Ins1 cells which were either wild-type (WT), ΔTBC1D23, or the latter rescued with TBC1D23-GFP stably expressed under a cumate-inducible promoter as indicated. (B) The intensities of the bands in (a) were quantified and normalised to wild-type (shown is mean and standard error of the mean (SEM)); number of biological replicates is 6 (HEK) or 3 (INS-1). (C) Confocal micrographs of WT or ΔTBC1D23 INS-1 cells stained for endogenous CPD and GM130 (Golgi marker). Scale bars: 10 µm. (D) Confocal micrographs of ΔTBC1D23 INS-1 cells transiently expressing mouse TBC1D23-GFP, and labelled for the GFP, and endogenous CPD and GM130. Scale bar: 10 µm. (E) Immunoblots of the indicated 293 cell lines as in (a) but with the addition of 293 cells expressing TBC1D23-GFP(1-558), and with quantification as in (b) showing mean and SEM of two independent replicates. (F) Immunoblots of whole cell lysates from wild-type 293 cells (WT) or the indicated mutants, some of which expressed TBC1D23-GFP from a cumate-inducible promotor. CPD was levels were quantified as in (b) for blots from three independent clones of ΔFAM21A/C, and ΔTBC1D23;ΔFAM21A/C, with mean values and SEM shown. CPD is destablised by removal of TBC1D23 but removal of FAM21A/C. Source data for Figs 2B, 2E and 2F are in data S2.

The C-terminal domain of TBC1D23 has been previously reported to bind to the FAM21 subunit of the WASH complex (*12*, *13*). However, removal of FAM21 from 293T cells did not affect the steady state levels of CPD (Fig. 2F). This indicates that the role of TBC1D23 in CPD traffic does not depend on the presence of FAM21. In contrast, deletion of the two golgins that bind TBC1D23 did reduce overall CPD levels as expected (fig. S1E). Taken together, these results show that TBC1D23 is required to maintain steady state levels of CPD in the TGN, and that performing this function involves an activity of TBC1D23 that is distinct from its ability to bind FAM21.

### Conserved residues in CPD mediate the interaction with TBC1D23

To gain more understanding of the interaction between TBC1D23 and CPD, we mapped the region of the CPD tail that is required for binding and found that a C-terminal 16 residue region is necessary and sufficient (Fig. 3, A and B). The residues Leu^1375^ and Tyr^1376^ are particularly important, with upstream conserved acidic residues also contributing which becomes more apparent when several are mutated (Fig. 3, B and C). NMR analysis of the CPD tail showed that the presence of TBC1D23 affected the residues in the C-terminal region around the acidic TLY motif, confirming it binding to this region in solution (fig. S2).

**Fig. 3.**
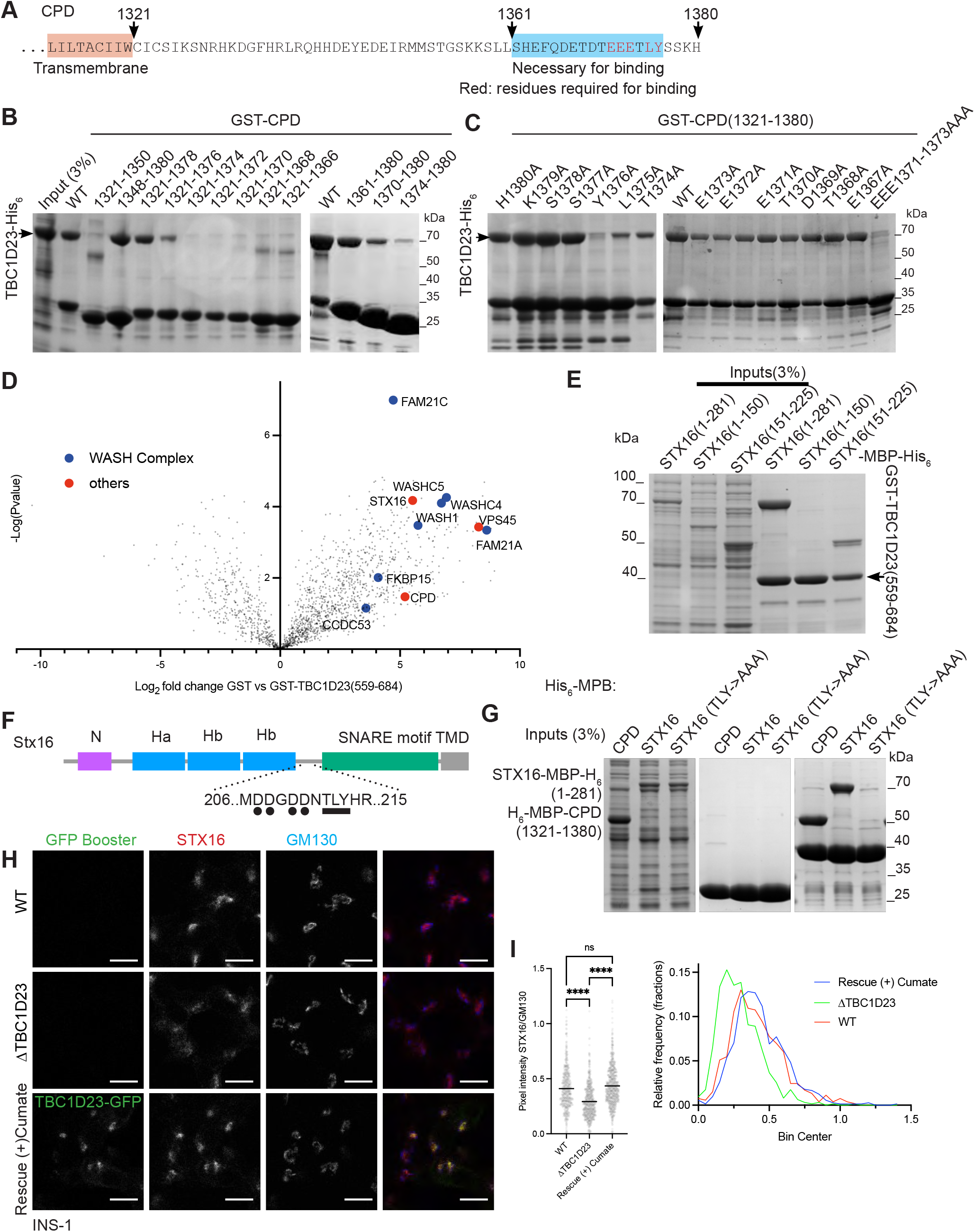
TBC1D23 binds to C-terminal conserved residues of the cytoplasmic tail of CPD. (A) The cytoplasmic tail of human CPD. The region highlighted in blue is sufficient for binding; residues in red are necessary. (B) Coomassie-stained gel testing which part of CPD is sufficient and required for binding to TBC1D23. GST-CPD constructs were immobilised on beads and incubated with a bacterial lysate containing TBC1D23-His_6_. (C) Coomassie-stained gel testing CPD mutants where the indicated residues were mutated to alanine. GST-CPD constructs were immobilised on beads and incubated with a bacterial lysate containing TBC1D23-His_6_. (D) Volcano plot of the mass spectrometry analysis from affinity chromatography of HEK-293T cell lysates using bacterially expressed GST-TBC1D23(559-684) or the negative control GST. The plot compares the spectral intensities from proteins bound to each bait, using data from three independent experiments. Values are in data S1. (E) Coomassie-stained gel showing that the C-terminal vesicle binding domain of TBC1D23 interacts with the syntaxin-16 using residues 151-225. Bacterially GST-TBC1D23(559-684) was immobilised beads and incubated with a bacterial lysate containing parts of syntaxin-16 tagged to MBP-His_6_. The numbering corresponds to the sequence of UniProt ID O14662-2 (F) Cartoon of syntaxin-16 showing the location of the key structural features and the acidic TLY motif related to that in the tail of CPD. (G) Coomassie-stained gel showing that residues T191, L192, Y193 of syntaxin-16 are required for binding to TBC1D23. GST-TBC1D23(559-684) was immobilised beads and incubated with a bacterial lysate His^6^-MBP-CPD(1321-1380), STX16(1-281)-MBP-His_6_ or STX16(1-281A, T191A, L192A, Y193)-MBP-His_6_. GST was used as a negative control. (H) Confocal micrographs of INS-1 cells (WT, ΔTBC1D23, and ΔTBC1D23 stably expressing TBC1D23-GFP under a cumate promoter in the presence of cumate for 24-36 hours). Cells were fixed, permeabilised and stained using GFP-booster and endogenous syntaxin-16 and GM130. (I) (left) Scatter plot showing the ratio of the Golgi-area fluorescence intensity of the syntaxin-16 channel over the GM130 channel. Golgi-area were automatically detected using a homemade ImageJ macro (see material and methods). The black horizontal bar is the mean. For WT, n = 530, for ΔTBC1D23, n = 628, for the stable rescues, n = 725. Statistics: ordinary one-way ANOVA followed by a Sidak’s multiple comparison tests with a single pooled variance. ****: P<0.0001, ns: not significant. (right) ratios were plotted as frequency distributions with a bin width of 0.05 (see material and methods). Source data is in data S2.

### TBC1D23 binds to the cytoplasmic domain of multiple cargoes of endosome-derived vesicles

The finding that TBC1D23 can bind directly to the cytoplasmic tail of CPD, a protein that is known to recycle from endosomes back to the Golgi, raised the possibility that this interaction could mediate the capture of endosome-derived vesicles by TBC1D23. However, the capture of such vesicles by TBC1D23 ectopically relocated to mitochondria still occurred in cells from which the CPD gene had been deleted, indicating that binding to CPD is not sufficient to account for vesicle capture (fig. S3, A to C). To find further factors that might contribute to vesicle capture in addition to CPD we used GST-TBC1D23 (559-684) for affinity chromatography of HEK-293T cell lysates. As expected, we identified CPD and subunits of the WASH complex amongst the interacting proteins (Fig. 3D). Also enriched was the SNARE syntaxin-16, a known cargo of endosome to Golgi carriers, along with one of its interacting partners, the SM protein VPS45 (*24*). We confirmed that syntaxin-16 is present in vesicles captured by either TBC1D23 or golgin-97 when they are relocated to mitochondria in HeLa cells (fig. S3, D and E). The syntaxin-16 cytoplasmic domain bound directly to the TBC1D23 C-terminal domain in vitro, and residues 151-225 of syntaxin-16 are both necessary and sufficient for the interaction (Fig. 3E). CPD binds to TBC1D23 via the sequence EEETLY, and interestingly syntaxin-16 contains a similar sequence: _209_DDNTLY in the exposed linker between the SNARE domain and the Habc domain (Fig. 3F). Mutating the TLY sequence in syntaxin-16 to AAA abolished its binding to TBC1D23 (Fig. 3G). While the steady state-levels of syntaxin-16 were not detectably altered in ΔTBC1D23 cells (fig. S3F), the Golgi-localised pool of the protein was reproducibly reduced, and this could be rescued by expression of TBC1D23-GFP (Fig. 3, H and I, and fig. S3G). Taken together, our results show that TBC1D23 binds to the cytoplasmic tail of at least two proteins in the vesicles that it captures and thus raise the possibility that this capture could occur via direct and specific interaction with a subset of vesicle cargoes.

### Structural basis of the interaction between TBC1D23 and syntaxin-16

To understand how the C-terminal domain of TBC1D23 recognises the tails of vesicle cargo we used a combination of X-ray crystallography and biochemistry. Peptides corresponding to the relevant regions of syntaxin-16 and CPD showed robust enthalpic-dominated binding to the C-terminal domain by Isothermal titration calorimetry (ITC) with affinities of ∼1.4 and 10.6 μM respectively and a stoichiometry of 1:1 (Fig. 4A). Crystallisation of the C-terminal domain in the presence of the syntaxin-16 peptide yielded crystals which diffracted to 3.3 Å resolution and were solved by molecular replacement using the previously published apo structure (6JM5) in space group P6_2_22 (Table S1).

**Fig. 4.**
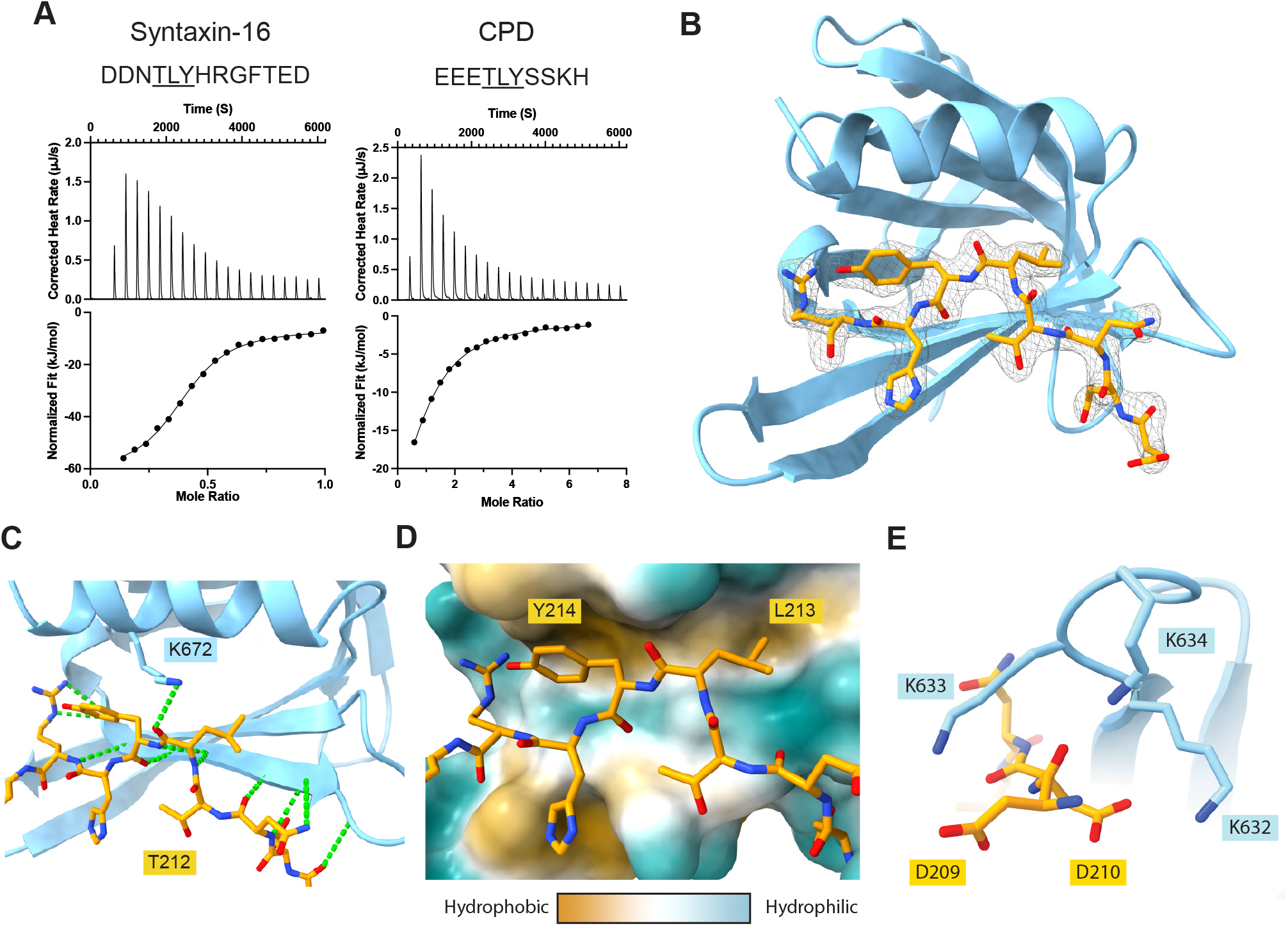
Structure of TBC1D23 C-terminal domain bound to a syntaxin-16 peptide. (A) ITC trace and fitted curve of the indicated peptides from syntaxin-16 (left) and CPD (right) binding to TBC1D23 C-terminal domain. (B) Structure of the TBC123 C-terminal domain (sky blue) and syntaxin-16 209-217 (gold) with the corresponding electron density map of the latter. (C) Close up of the hydrogen bonding network between syntaxin-16 (gold) and TBC1D23 (sky blue). K672 hydrogen bonds to the backbone of syntaxin-16. Hydrogen bonds in green. (D) Hydrophobicity surface plot of TBC1D23 with packing of syntaxin-16 (gold) into two hydrophobic pockets. (E) Close up of aspartic acids 209 and 210 in syntaxin-16 (gold) interacting with lysines 632-634 in TBC1D23 (sky blue).

Four C-terminal domains were present within the asymmetric unit (fig. S4A-C). Two of the domains dimerised via strand exchange in the same manner as that seen in the peptide-free structure with reciprocal C-terminal tails (VLDALES) inserting between strand 5 and the helix 2 of the other domain, but with no sign of the peptide (*13*). In the third domain, the electron density was unclear, but for the final domain density corresponding to a single syntaxin-16 peptide was visible (fig S4D). This peptide density was in the same position that was occupied by the cross-dimerising, C-terminal tail in the first two molecules of the asymmetric unit.

Given the above results, we generated a version of the C-terminal domain lacking the last seven residues (VLDALES). Crystals of this truncated domain in the presence of the syntaxin-16 peptide diffracted to higher resolution (2.2 Å) and were solved in space group P6_5_22 again by molecular replacement with unliganded 6JM5 (Table S1). The asymmetric unit contained two copies of the domain with both having density corresponding unambiguously to the syntaxin-16 peptide in the previously proposed binding site (fig. S4E). The previously observed strand-exchanged dimer was absent with the homodimeric C-terminal tail-mediated interface being replaced by the peptide. Thus, the structure elucidates how the syntaxin-16 peptide binds to the C-terminal domain of TBC1D23 (Fig. 4B). The peptide extends TBC1D23’s β sheet by adopting a twisted β augmentation mode of interaction with strand 5, and buries ∼670 Å^2^ of the C-terminal domain’s solvent accessible surface area, a similar sized interface to those seen previously for other trafficking motif-binding proteins that have a similar K_D_ of interaction (*25*, *26*). The Thr^212^ in the acidic TLY motif of syntaxin-16 packs its methyl group against the alkyl chain of Lys628 which orients the hydroxyl group towards the solvent to stabilise a somewhat strained backbone conformation (Fig 4C). This allows the sidechains of the leucine and tyrosine residues to be splayed apart and simultaneously buried into neighbouring hydrophobic pockets in the domain (Fig. 4D). As a result, the acidic residues Asp^209^ and Asp^210^ of syntaxin-16 are in close proximity to three basic residues (lysines 632-634) in the C-terminal domain of TBC1D23, which would allow a direct electrostatic interaction (Fig. 4E).

### Mutants in the TBC1D23 C-terminal domain and the syntaxin-16 peptide perturb binding

The structure indicates that recognition of the acidic cluster and the TLY triad are the critical factors in driving peptide binding and specificity. To validate this, a series of point mutations was designed in the C-terminal domain and initially assessed for folding by yield and circular dichroism. Two of the residues that form parts of the hydrophobic pockets that bind the L and Y of the peptide (I^629^ and I^639^), are also part of the domain’s core hydrophobic residues and their mutation caused misfolding (I629S, I639S). The remaining mutants were assayed for binding to the syntaxin-16 peptide by ITC (Fig. 5A-C). K^672^ of TBC1D23 hydrogen bonds via its amine group to the carboxyl peptide bond of L^213^ in the TLY in syntaxin-16 (Fig. 4C); mutation to alanine (K672A) weakened binding ∼20 fold (K_D_∼30 μm). Mutation of Val^626^, which packs against the aromatic ring of reside Y^214^, to aspartate (V626D) abolishes peptide binding. The binding pocket in TBC1D23 for Y^214^ was obstructed by mutating one of its lining residues (I675W), and again this reduced binding to below detectable levels (K_D_ >300µM). Finally, mutation to alanine of lysines 632-634, which are adjacent to the TLY-binding hydrophobic grove, also abolished peptide binding, indicating that their interaction with the acidic cluster in syntaxin-16 is important. Thus, mutations which either remove interactions with the peptide, or protrude into the binding pocket strongly affect peptide binding confirming that the interactions seen in the crystal also occur in solution.

**Fig. 5.**
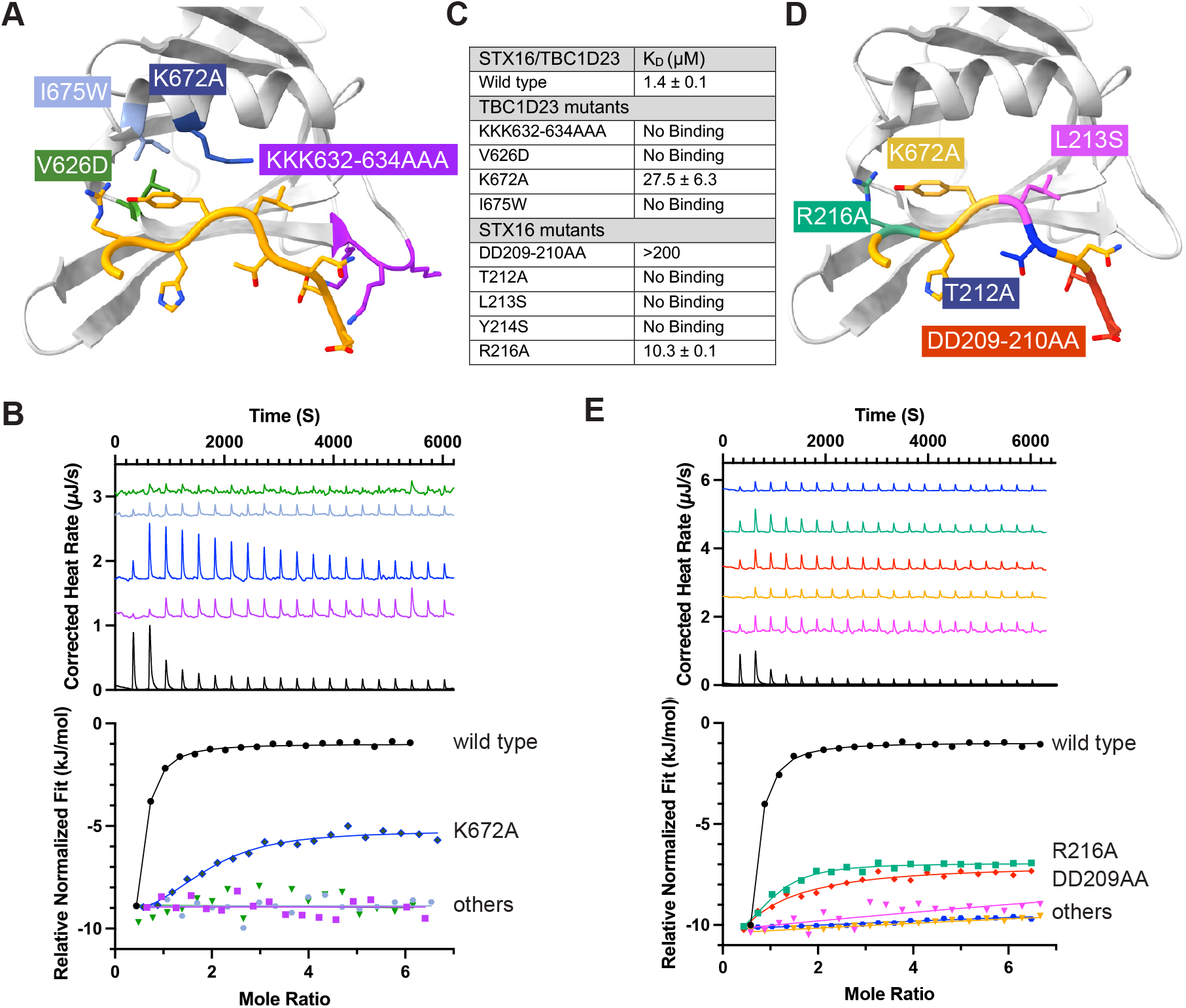
Mutational analysis of the interaction between TBC1D23 and syntaxin-16. (A) Schematic view of the mutations in TBC1D23 that were tested by ITC. (B) Isothermal titration calorimetry of relative binding of syntaxin-16 peptide to the TBC1D23 C-terminal domain mutants. Mutations are indicated by the colour coding in (A), with wild type in black. (C) Summary of ITC results. (D) Schematic view of the mutations in the syntaxin-16 peptide that were tested by ITC. (E) Isothermal titration calorimetry of relative binding of the mutant syntaxin-16 peptides to the TBC1D23 C-terminal domain. Mutations are indicated by the colour coding in (D), with wild type in black.

Mutations were also made in the syntaxin-16 peptide to further validate the interactions observed in the crystal structure (Fig. 5D). ITC assays confirmed a role in binding for all three of the residues in the acidic TLY motif and for the two adjacent acidic residues, with R^216^ making a smaller contribution (Fig. 5C-E). The acidic TLY motif is absolutely conserved between syntaxin-16 and CPD, and so to determine if conservative changes in the motif can be tolerated, we tested variants of these three residues (fig. S4F). Mutation of T^212^, which holds the leucine and tyrosine in a strained position to allow engagement with their pockets, to serine (T212S) reduced binding ∼20 fold to K_D_∼30 µM. Changing L^213^ to the smaller alanine (L213A) or the larger phenylalanine (L213F) both caused a reduction of binding of >100 fold, consistent with the side-chain fitting into a size-specific hydrophobic pocket in the structure, i.e. a medium sized hydrophobic residue is required at this position. Finally, Y^214^ is more tolerant of conservative mutation with Y214W and Y214F peptides having K_D_s of ∼7 µM and ∼15 µM respectively (fig. S4F).

### Use of the C-terminal-domain binding motif to identify further binding partners

Although both syntaxin-16 and CPD bind to the C-terminal domain through a closely related motif, the only functional property that they share is the fact that they are both in vesicles that recycle between endosomes and Golgi and can be captured by TBC1D23. We thus wondered if there were further proteins in these vesicles that also have an acidic TLY motif that can bind TBC1D23. Peptide motifs that are recognised by proteins are rarely invariant, and so other versions of the motif are likely to be functional. From the above mutagenesis experiments and the nature of the interactions seen in the structure, we chose [DE]_>=3/5_[TS][LIV][YFW] as being a loose but plausible definition of the core binding motif for the TBC1D23 C-terminal domain and used this to search the predicted cytoplasmic tails of all membrane proteins in the human proteome. In addition, we looked for membrane proteins amongst the interactors reported for TBC1D23 in the BioPlex protein interaction screen (*27*). The latter approach identified four membrane proteins, three of which are reported to be in the endosome/Golgi system, and one of unknown location. The former approach identified 19 membrane proteins of which 10 are reported to be endosomal or Golgi, with two proteins found by both approaches (Table S2). We thus tested the ability of the TBC1D23 C-terminal domain to bind to peptides corresponding to the TLY-like region from several of these proteins (Fig. 6A-C and fig. S5).

**Fig. 6.**
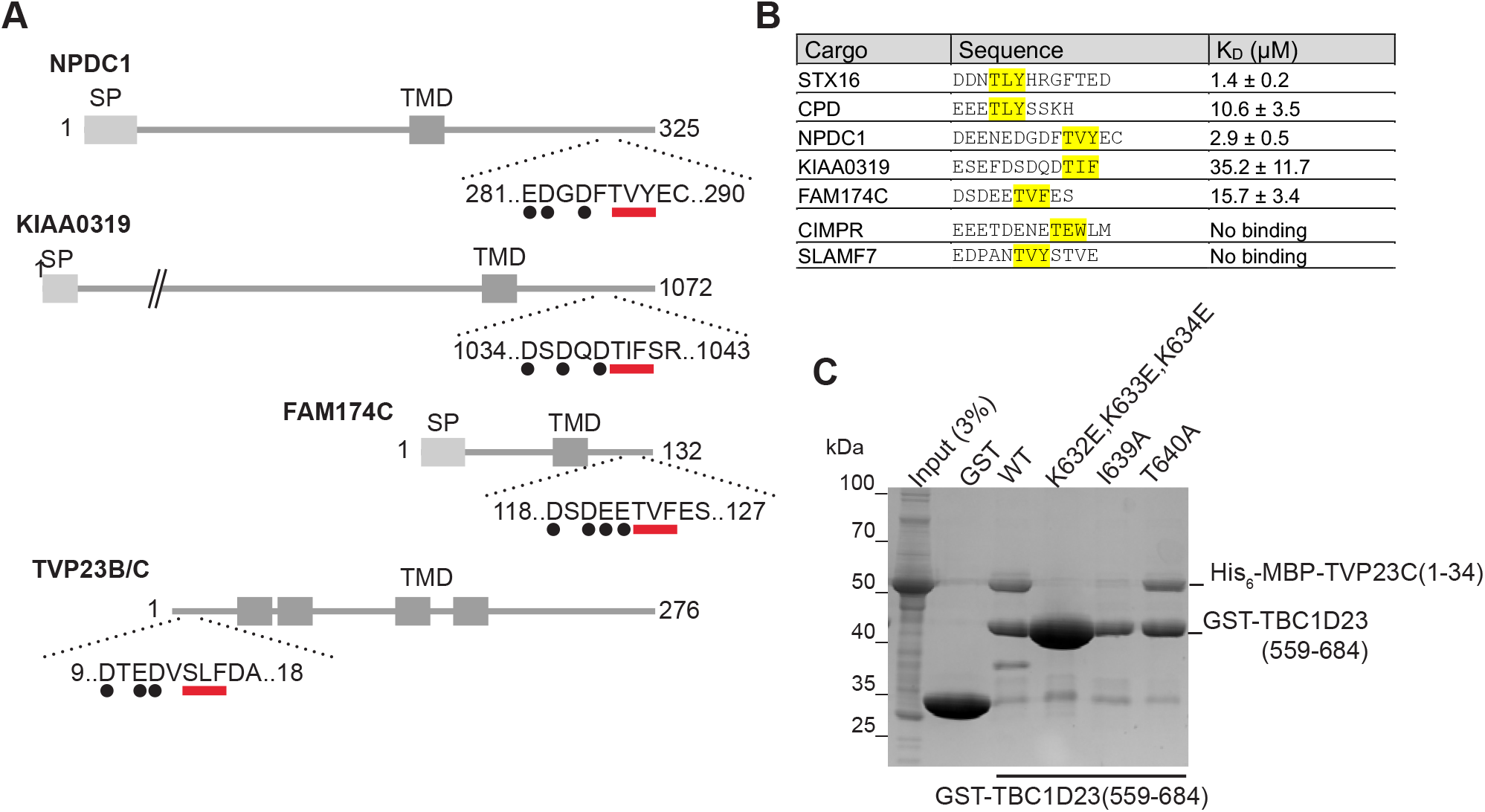
Identification of membrane proteins containing a TBC1D23-binding acidic TLY motif. (A) Schematic of four human proteins that have an acidic TLY motif in the cytoplasmic tails and which have been reported to recycle from endosomes to the Golgi, or be localised to the Golgi. (B) Binding affinities of the indicated peptides covering to the acidic TLY motifs from the proteins shown in (A), as determined by ITC (fig. S5). (C) Coomassie-stained gel showing the binding of the N-terminal cytoplasmic tail of TVP23C (His_6_-MBP-TVP23C(1-34) to beads coated with the C-terminal domain of TBC1D23 fused to GST. Mutations in the acidic TLY motif disrupt the interaction.

NPDC1 is a largely uncharacterised protein suggested to play a role in neuropeptide secretion and dense core vesicle traffic (*28*, *29*), and the peptide from its tail has an affinity for TBC1D23 similar to that of syntaxin-16 at ∼3 µM (fig. S5). KIAA0319 is a neuronal protein linked to dyslexia, and although its function remains unknown, it has been reported to recycle from the surface back to the Golgi region (*30*). The peptide from KIAA0319 peptide showed weaker, but detectable binding with K_D_∼ 35 µM (fig. S5). FAM174A and its paralogues FAM174B and FAM174C are small membrane proteins of unknown function that have been reported to be in the Golgi (*31*). All have a TLY-like motif and the peptide from FAM174C bound with an affinity of 16 µM. Finally, TVP23B/C is a small polytopic membrane protein known to recycle between endosomes and Golgi (*32*, *33*), and its 34 residue cytoplasmic tail bound to the TBC1D23 C-terminal domain with this interaction disrupted by mutating the three lysine residues in TBC1D23 that we have shown to be required for binding CPD and syntaxin-16 (Fig. 6C). In contrast, no binding was seen with peptides from CIMPR, which also traffics between Golgi and endosomes and has an acidic region in its tail but no TLY-like motif, and SLAMF7 which has a TLY-like motif but smaller acidic cluster that contains a proline (Fig. 6B). Thus, multiple proteins that recycle between endosomes and Golgi have sequences in their cytoplasmic tails that can bind directly to the C-terminal domain of TBC1D23.

### Residues in TBC1D23 that are required for binding the acidic TLY motif are also required for vesicle capture in vivo

The presence in endosome-to-Golgi vesicles of diverse proteins with a TLY-like motif in their cytoplasmic tails raises the possibility that TBC1D23 can capture these vesicles by binding to the tails protruding from the vesicle. TBC1D23 is sufficient to capture vesicles when relocated to mitochondria, and so we tested the effect on this capture of mutations that disrupt peptide binding in vitro. Relocation of vesicles by the constructs was quantified with BioID proximity biotinylation by using a promiscuous biotin ligase fused to TBC1D23 along with the mitochondrial targeting signal (Fig. 7A). When wild-type TBC1D23 was expressed on mitochondria, vesicle cargo was efficiently biotinylated as expected (Fig 7B). However, mutation of the residues that disrupt peptide binding in vitro resulted in a near complete loss of biotinylation of vesicle cargo proteins indicating that vesicle capture was also disrupted. This is not simply due to the vesicle being docked by a different mechanism, as immunofluorescence demonstrated that these cargo proteins were no longer accumulating on the TBC1D23-coated mitochondria (Fig 7C). Taken together, these results demonstrate that the same part of TBC1D23 that binds to the tails of vesicle cargo in vitro is required for vesicle capture in vivo, consistent with a model in which TBC1D23 can capture the incoming vesicle by recognising its cargo proteins.

**Fig. 7.**
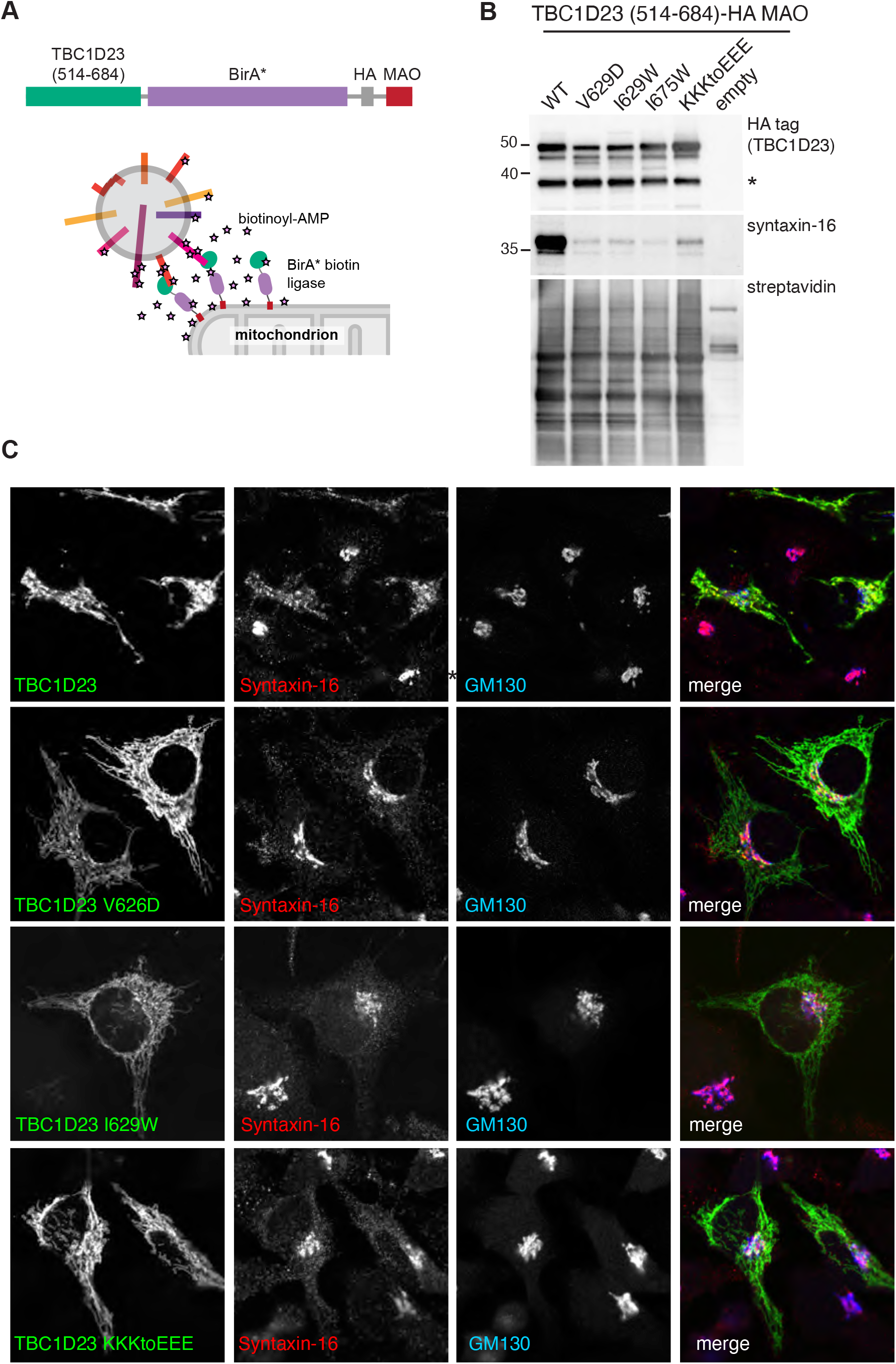
Residues in TBC1D23 required for peptide binding in vitro are also required for vesicle capture in vivo. (A) Ectopic relocation and biotinylation: a chimera comprising the TBC1D23 C-terminal domain attached to the BirA* promiscuous biotin ligase and the mitochondrial targeting signal from monoamine oxidase. (B) Immunoblots of total cell extracts from HEK293T cells transfected with the indicated variants of the mitochondrially targeted C-terminal domain fused to BirA*. Mutation of key residues does not affect total biotinylation or the stability of the chimera, but greatly reduces biotinylation of syntaxin-16 indicating loss of vesicle capture. The lower band (asterisk) indicates some clipping of the chimera apparently between the TBC1D23 part and BirA*. (C) Immunofluorescence of cells expressing the indicated forms of mitochondrial full-length TBC1D23 and immunolabelled for the TBC1D23 chimera (HA tag), syntaxin-16, and the Golgi marker GM130. The mutations in TBC1D23 that disrupt peptide binding in vitro greatly reduce mitochondrial accumulation of syntaxin-16 containing vesicles.

## Discussion

The C-terminal domain of TBC1D23 was known to be sufficient to capture one or more of the classes of carrier that mediate transport of membrane proteins from endosomes to Golgi (*12*, *13*). Our search for binding partners of this domain has now revealed that it can bind directly to the cytoplasmic tails of several of the cargo proteins that are present in these vesicles via a conserved acidic TLY motif. This interaction provides a mechanism by which TBC1D23 displayed on the ends of golgin-97 and golgin-245 can recognise specific vesicles and thus tether them to the trans Golgi prior to subsequent SNARE engagement and vesicle fusion. For both CPD and syntaxin-16, their predicted structures indicate that the acidic TLY motif is likely to be located at some distance from the TMD and hence the vesicle membrane. The unstructured tail of CPD or the α-helical bundle of syntaxin-16 are predicted to place the motif ∼15 nm away from the membrane which would allow recognition to be unimpeded by the vesicle surface.

The direct binding of a tethering factor to the cargo carried by a vesicle has not been previously proposed as a mechanism for vesicle capture but it would certainly ensure that only a specific set of vesicles is recognised (34). Like all such tethering interactions, the mechanism requires that the tether binds only to the vesicle and not to the organelle from which the cargo-laden vesicle budded. It is possible to imagine several ways in which this could be achieved. TBC1D23 is primarily localised at the Golgi as determined by both immunofluorescence and organelle fractionation, indicating that the interaction of its N-terminal domain with golgin-97 and golgin-245 is the dominant factor in determining its localisation in vivo (*12*, *33*, *35*). In addition, the golgins are homodimers and so each golgin is likely to display two copies of TBC1D23 which would increase the avidity of the interaction with the tails in the vesicle over that of a monomer of TBC1D23 that was free in the cytoplasm. Indeed, it may be that interaction between TBC1D23 and the cargo proteins is only strong enough to capture a vesicle when an array of TBC1D23 presented on multiple golgins can interact with multiple TLY-containing proteins that are concentrated in a single vesicle and hence the avidity of the interaction is increased even further.

Another mechanism that could account for TBC1D23 only recognising cargo proteins that are present in vesicles is that the acidic TLY motif in the cargo proteins could be masked in some way when the cargo is in endosomes and other compartments. This could occur if the coat proteins that sort them into the retrograde pathway rapidly sequester them into forming carriers, and that the interaction with the coat prevents recognition of the acidic TLY motif by TBC1D23 until the carrier has budded and uncoated. The identity of the coat that makes the carriers captured by TBC1D23 is as yet unresolved as there are multiple routes back from endosomes to the Golgi, with both clathrin coats with the AP-1 adaptor and also sorting nexins with retromer proposed to be involved in generating carriers (*19*, *36*). Nonetheless, it is worth noting that the acidic TLY motif includes a stretch of acidic residues, and acidic clusters can act as sorting signals for packaging into clathrin/AP-1 coated carriers (*37*, *38*). Both CPD and syntaxin-16 are known to be enriched in AP-1-dependent clathrin-coated vesicles, although whether such vesicles are delivering cargo to or from the trans Golgi also remains to be resolved (*39*).

There are known to be multiple classes of vesicle arriving at the Golgi from endosomes, and although TBC1D23 appears to responsible for the capture of at least some of these, the mechanism by which the others are captured is unknown. The golgin GCC88 captures at least one such set of vesicles even though it does not bind TBC1D23, but how this is achieved is still unclear (*9*, *12*). This multiplicity of endosome to Golgi routes with only some carriers being captured by TBC1D23 is consistent with human mutation of TBC1D23 resulting primarily in neurodevelopmental defects despite being widely expressed in most tissues (*35*, *40*). It may be that when TBC1D23-dependent tethering is lost, other capture mechanisms can partially compensate, or some cargo proteins can return to the Golgi by some of the alternative routes.

The enrichment of a particular set of proteins in each specific type of transport vesicle is only one type distinguishing feature that would allow the specific recognition necessary to ensure the fidelity of intracellular traffic. For most trafficking steps, vesicle recognition remains poorly understood. In some cases Arf or Rab GTPases are believed to play a role, although they too are likely to be present on donor organelles resulting in the same issues of vesicle versus donor organelle identity (*4*, *34*, *41*). Our finding that a tethering factor can recognise a set of cargo proteins in the carriers that it captures not only sheds light on endosome to Golgi traffic but could also illuminate other membrane trafficking routes where the mechanism of vesicle recognition remains to be discovered.

## Materials and Methods

### Plasmids

Details of the plasmids used in this report, together with the cloning methods used to generate are provided in data S3. Note that the cDNA for TBC1D23 used throughout is from mouse (Q8K0F1) with 684 residues (corresponding to the 684-residue human isoform (UniProt Q9NU8-2)), and residues are numbered accordingly.

### Antibodies

Full lists of primary and secondary antibodies used for western blotting and immunofluorescence are provided in data S3.

### Mammalian cell culture

The full list of cell lines used are in data S3. HeLa (ATCC), Human embryonic kidney 293 T (HEK293T, ATCC, CRL-3216), and HEK 293 Flp-In™ T-REx™ 293 cells stably expressing Cas9 (referred to throughout the paper as HEK-293) were cultured in Dulbecco’s modified Eagle’s medium Glutamax (DMEM; Gibco) supplemented with 10% foetal calf serum (FCS) and penicillin/streptomycin at 37°C and 5% CO_2_. The insulinoma INS-1-derived 832/13 rat cell line was obtained from Christopher Newgard (Duke University School of Medicine) via Michael Ailion (University of Washington, Seattle) (referred to throughout the paper as INS-1, (*42*)). INS-1 cells were grown with RPMI-1640 Glutamax (GIBCO), supplemented with 10% FCS, 1 mM sodium pyruvate, 10 mM HEPES, and penicillin/streptomycin at 37°C and 5% CO_2_. For transient transections, unless noted, we used polyethylenimine (PEI; Polyscience, 24765) dissolved in PBS to 1 mg/mL. The ratio of PEI (µL) to DNA (µg) used was 3:1. PEI was dissolved in Opti-Mem (Gibco) and incubated at room temperature for 5 minutes. DNA was added and incubated for another 20 minutes at room temperature before dropwise addition onto cells which had been seeded the day before. Cells were free of mycoplasma as determined by routine testing using MycoAlert (Lonza).

### Generation of knock-out cells using CRISPR-Cas9

Precise information on the cell lines generated as well as gRNA sequences is provided in data S3. ΔTBC1D23 (in HEK293 Flp-In™ T-REx™ 293 cell line stably expressing Cas9) cells were generated as follows: cells grown to ∼70% confluence in 6-well plates and co-transfected with a gRNA to TBC1D23 (*12*) and pmaxGFP using Fugene6 (Promega) according to manufacturer’s instruction. 48 hours later, cells were trypsinised, and single GFP positive cells were sorted into each well of 96-well plates using a FACS sorter. Single colonies that grew were expanded and analysed by immunoblotting.

All other knockouts made in HEK293T, HEK293 Flp-In™ T-REx™ 293 stably expressing Cas9, and INS-1 cells were generated as follows: gRNA with high efficiency and high specificity scores were chosen using the UCSC genome browser and CRISPOR (*43*). To improve the knockout efficiency, each gene was targeted with two gRNA, one targeting an early exon and the second targeting a late exon, so a large part of the coding region could be removed (data S3). All gRNAs were cloned onto the pX459 vector; which also expresses Cas9 and a puromycin resistance gene (*44*). Cells were grown to ∼70% confluence in 6-well plates and transfected with a mixture of 6 μL of PEI and equal amount of each vector coding for gRNAs for a total of 2 μg of DNA in 100 μL of Opti-MEM. 48 hours later, the cells were trypsinised and replated in complete medium containing 1.5 μg/mL puromycin. 48 to 72h hours later, cells were diluted to 1 cell per two wells in three 96-well plates and grown in complete medium. Cloned that grew were expanded and screened by immunoblotting or immunofluorescence. When possible, several clones were assayed (data S3).

For rescue of knockout cell lines, the relevant genes were cloned in to a piggyBac vector containing a puromycin resistance gene and the protein of interest expressed under a cumate-inducible promoter. Cells grown to ∼70% confluence in 10 cm plates were co-transfected with the plasmid coding for the rescue constructs and the plasmid coding the piggyBac transposase at 5:1 molar ratio using 6 µg of DNA, 18 µL of PEI in 1 mL Opti-MEM. 48 hours after transfection cells were trypsinised and replated in selection media containing 1.5 µg/mL Puromycin. The media was changed every 48-72 h while keeping the puromycin selection until confluence was reached (usually 10 days), and expression of the rescue construct was verified by western blotting and immunofluorescence. Rescue lines were maintained as a polyclonal population.

### Immunofluorescence

Cells were transfected in 24-well plates (300 ng of DNA 1 μL of PEI in 50uL Opti-MEM) or in 6-well plates (1 µg of DNA, 3µL of PEI in 100µL of Opti-MEM). The next day, cells were dissociated using trypsin and seeded onto PTFE-coated multiwell slides (Hendley-Essex). ∼36 to 48h post-transfection cells were fixed with 4% paraformaldehyde in PBS (10 minutes), permeabilised in 0.5% (v/v) Triton X-100 in PBS (5 minutes), and blocked for one hour in PBS containing 20% FCS and 0.25% Tween-20. Primary and secondary antibodies were applied sequentially in blocking buffer for 1h at room temperature. After washing, cells were mounted using ProLong Gold antifade mountant (Thermo Fisher Scientific). Slides were imaged using a Leica TCS SP8 laser scanning confocal microscope and a 63X lens.

### Quantification of fluorescent micrographs

For each condition in each independent replicate, 3 multi-channel fluorescence micrographs at 1024x1024 resolution were acquired. The images were then automatically processed using a custom ImageJ macro to measure the intensity in all channels for regions of interest (ROI) defined by segmenting the second channel (Golgi marker GM130). In short, the second channel was processed with a rolling ball and a median filter. A threshold based on the mean and standard deviation of the image allowed to separate the background and foreground while a filter on minimum area remove spurious detection. ROI were enlarged in order to include the nearby region. Finally, the macro reports in a table the area of the ROI and the mean and maximum value of each channel. Segmentation can be visually assessed by inspecting overlays saved in a TIF file. For each condition over 100 individual cells were quantified. For each ROI, the mean fluorescence intensity of the protein of interest was divided by the mean fluorescence intensity of the GM130 channel. The data of the ratio of mean fluorescence intensities were then plotted as frequency distributions using Prism 9 with a bin width of 0.05. For each marker, the experiment was repeated twice with similar results.

### Determination of protein levels by immunoblotting

HEK293 or INS-1 cells were seeded in 10-cm plates, and where appropriate, the media was supplemented with 1X cumate (System Biosciences QM159A-1) to induce expression of the rescue constructs. After 24 to 36 hours, cells were resuspended by scraping, washed twice with ice-cold PBS, and pelleted by a 5-minute 500 x g spin at 4°C. Cells were resuspended in lysis buffer (50 mM Tris, pH 7.4, 150 mM NaCl, 1 mM EDTA, 1% Triton X-100, and protease inhibitor cocktail (cOmplete, Roche)), and incubated on ice for 5 minutes. Lysates were clarified at 17,000 x g for five minutes at 4°C, and the total protein concentration was determined using the BCA assay (Thermo Fisher Scientific 23227). Clarified lysates were then supplemented with 4x NuPage LDS sample buffer containing 100 mM DTT (Invitrogen, NP0007). Equal amount of total protein was separated on SDS PAGE gels (Invitrogen, XP04205) and analysed by immunoblotting.

### Immunoblotting

Protein samples in 1x NuPage LDS sample buffer containing 25 mM DTT were boiled at 95°C, loaded on to SDS PAGE gels (Invitrogen, XP04205) and transferred to nitrocellulose membranes. Membranes were blocked in 5% (w/v) milk in PBS-T (PBS with 0.1% [v/v] Tween-20) for 1 hour, incubated overnight at 4°C with primary antibody in the same blocking solution, washed three times with PBS-T for 5 minutes, incubated with HRP-conjugated secondary antibody in 5% (w/v) milk in PBS-T for 1 hour and, washed three times with PBS-T for 5 minutes. Where indicated a primary anti-6xHis antibody crosslinked to HRP was used. Blots stained with HRP-conjugated secondary antibodies were imaged using BioRad ChemiDoc imagers with SuperSignal West Pico PLUS (Thermo Fisher Scientific, 34577). The “gels” analysis tool in ImageJ was used for the quantification of blots. For each blot, raw data was normalised to the WT band. The raw data can be found in the source file.

### *In vitro* binding assays using recombinant proteins

Recombinant proteins for binding assays were expressed as follows: plasmids were transformed into E. coli BL21-CodonPlus (DE3)-RIL (Agilent, 230245). From an overnight starter culture, cells were grown in 2 x TY medium containing 100 µg/mL ampicillin (or 50 µg/mL kanamycin when appropriate) and 34 µg/mL chloramphenicol at 37°C in a shaking incubator. When the culture reached OD_600_ = 0.6-0.8, the temperature was lowered to 16°C, protein expression was induced with 100 µM of IPTG, and incubated overnight. Bacteria cells were harvested by centrifugation at 4,000 x *g* at 4°C for 15 minutes and were mechanically resuspended on ice in lysis buffer containing 50 mM Tris, pH 7.4, 150 mM NaCl, 1 mM EDTA, 5 mM 2-mercaptoethanol, 1% Triton X-100, and protease inhibitor cocktail (cOmplete, Roche). Cells were lysed by sonication and the lysates were clarified by centrifugation at 20,000 x *g* at 4°C for 15 minutes. Clarified lysates were flash frozen in liquid nitrogen and stored at -80°C until needed.

For binding to beads, saturating amounts of clarified bacterial lysates containing GST-tagged baits were added to glutathione-Sepharose beads previously washed with lysis buffer (50 mM Tris, pH 7.4, 150 mM NaCl, 1 mM EDTA, 5 mM 2-mercaptoethanol, 1% Triton X-100) and incubated at 4°C for 1 hour on a tube roller. Beads were washed once with lysis buffer, once with lysis buffer supplemented with 500 mM NaCl, once again with lysis buffer, and incubated with clarified bacterial lysates containing the recombinant prey for 2 hours at 4°C on a rotator. Beads were washed 3 times with lysis buffer and eluted by boiling in lysis buffer supplemented with a 4x solution of NuPage LDS sample buffer containing 100 mM DTT. Boiled slurry was separated on SDS PAGE gels (Invitrogen) and analysed by Coomassie blue stain (Abcam, ab119211) or by immunoblot using an anti-6xHis HRP-conjugated antibody.

### TBC1D23 affinity chromatography from HEK293T cell lysate

GST-TBC1D23(1-513) and GST-TBC1D23(514-684) were expressed in the *E. coli* strain BL21-GOLD (DE3; Agilent Technologies). Bacteria were grown at 37°C to an OD600 of 0.7 and induced with 100 μM isopropyl β-D-1-thiogalactopyranoside (IPTG) overnight at 16°C. Cells were harvested by centrifugation, resuspended in lysis buffer (25mM Tris-HCl, pH 7.4, 150 mM NaCl, 1mM EDTA, 1% (v/v) Triton X-100, plus 1 EDTA-free complete protease tablet/50 ml and 1mM PMSF), dounce homogenised and sonicated on ice. The lysates were clarified by centrifugation at 12,000 x *g* for 15 min at 4°C and applied at saturating levels to glutathione-Sepharose beads. The beads were then incubated with cell lysates prepared from two confluent 175 cm^2^ flasks of HEK293T cells. HEK293T cells were collected by centrifugation at 500 x g for 3 min, washed once in ice-cold PBS and lysed in lysis buffer (25 mM Tris-HCl, pH 7.4, 150 mM NaCl, 1mM EDTA, 1% (v/v) Triton X-100 plus 1 cOmplete protease inhibitor tablet/50 ml and 1mM PMSF) at 4°C for 30 min followed by clarification by centrifugation at 17000 x g for 10 minutes. Lysates were pre-cleared on empty glutathione-Sepharose beads for 30 min prior to a two-hour incubation with TBC1D23-coated beads. Beads were washed extensively in lysis buffer and proteins eluted first in a high salt elution buffer (25mM Tris-HCl, 1.5M NaCl, 1mM EDTA) to release interacting proteins and then in SDS sample buffer to release the GST-fusion proteins. Proteins were precipitated from the high salt elution sample by chloroform/methanol precipitation and resuspended in SDS sample buffer containing 1 mM 2-mercaptoethanol. Eluates were separated by SDS-PAGE electrophoresis and analysed by mass spectrometry.

### Affinity chromatography of HEK293T cell lysate with GST-tail fusions

The approach used was as described previously (*45*), as follows. Clarified lysates from 450 mL 2xTY cultures containing bacteria expressing recombinant GST, GST-CPD(1321-1380), GST-CIMPR(2327-2491) and GST-TBC1D23(559-684) were thawed. For each GST-tagged bait, 100 μL of glutathione Sepharose 4B bead slurry (GE Healthcare) was used. Clarified bacterial lysates were added to the empty glutathione-Sepharose beads and incubated at 4°C for 1 hour on a roller. HEK-293T cells (from four confluent T175 flasks per GST-tagged bait) were collected by scraping, washed twice with PBS, and lysed with lysis buffer (50 mM Tris, pH 7.4, 150 mM NaCl, 1mM EDTA, 5 mM 2-mercaptoethanol, 1% Triton X-100, EDTA-free cOmplete protease inhibitor). The lysate was clarified by centrifugation for 5 minutes at 17,000 *x g* and pre-cleared with 100 μL of bead slurry for 1h at 4°C on a tube roller. Beads loaded with recombinant GST-tagged baits were washed once with ice-cold lysis buffer, once with lysis buffer supplemented with 500 mM NaCl, and once again with lysis buffer. Beads were incubated with the pre-cleared HEK-293T cell lysate for 2-4 hours on a roller at 4°C. Beads were washed twice with lysis buffer, transferred to 0.8 mL centrifuge columns (Pierce 89869B) and washed twice more. Columns were brought to room temperature and eluted 5 times with 100 µL of elution buffer (1.5 M NaCl in lysis buffer) by centrifugation at 100 x *g* for 1 minute; for the final elution the sample was centrifuged at 17,000 x *g* for 1 minute. Eluates were pooled together and concentrated to ∼75 μL using a centrifugal filter (Amicon Ultra 0.5 mL 3,000, Millipore UFC500324), supplemented with 25 µL of NuPage 4x LDS sample buffer (Invitrogen, NP0007) containing 100 mM DTT. 40% of the eluate was separated by SDS PAGE (Invitrogen, XP04202) and stained with InstantBlue Coomassie stain (Abcam, ab119211). Each lane was cut into 8 gel slices, transferred into a 96-well microtiter plate and subjected to mass spectrometry analysis.

### Mass spectrometry analysis of eluates from affinity chromatography

For samples prepared with GST-fusions to parts of TBC1D23, slices were destained with 50% v/v acetonitrile and 50 mM ammonium bicarbonate, reduced with 10 mM DTT, and alkylated with 55 mM iodoacetamide. After alkylation, proteins were digested with Trypsin (Promega, UK) overnight at 37 °C at an enzyme to protein ratio of 1:20. The resulting peptides were separated by nano-scale capillary LC-MS/MS using an Ultimate U3000 HPLC (ThermoScientific Dionex, San Jose, USA) to deliver a flow of approximately 300 nL/min. A C18 Acclaim PepMap100 5 µm, 100 µm x 20 mm nanoViper (ThermoScientific Dionex, San Jose, USA), trapped the peptides prior to separation on a 25 cm PicoCHIP nanospray column packed with Reprosil-PUR C18 AQ (New Objective Inc., Littleton, USA). Peptides were eluted with a 60-minute gradient of acetonitrile (2%v/v to 80%v/v). The analytical column outlet was directly interfaced via a nano-flow electrospray ionisation source, with a hybrid dual pressure linear ion trap mass spectrometer (Orbitrap Velos, ThermoScientific, San Jose, USA). Data-dependent analysis was carried out, using a resolution of 30,000 for the full MS spectrum, followed by ten MS/MS spectra in the linear ion trap. MS spectra were collected over a m/z range of 300–2000. MS/MS scans were collected using a threshold energy of 35 for collision induced dissociation. LC-MS/MS data were then searched against a protein database (UniProt KB) using the Mascot search engine programme (Matrix Science) (*46*). Database search parameters were set with a precursor tolerance of 5 ppm and a fragment ion mass tolerance of 0.8 Da. Two missed enzyme cleavages were allowed and variable modifications for oxidised methionine, carbamidomethyl cysteine, pyroglutamic acid, phosphorylated serine, threonine and tyrosine were included. MS/MS data were validated using the Scaffold programme (Proteome Software Inc.) (*47*).

For samples prepared with GST-fusions to tails of cargo proteins, gel slices (1-2 mm) were placed in 96-well microtiter plates and de-stained with 50% v/v acetonitrile and 50 mM ammonium bicarbonate, reduced with 10 mM DTT, and alkylated with 55 mM iodoacetamide. After alkylation, proteins were digested with 6 ng/µL trypsin (Promega, UK) overnight at 37 °C. The resulting peptides were extracted in 2% v/v formic acid, 2% v/v acetonitrile. The digests were separated by nano-scale capillary LC-MS/MS using an Ultimate U3000 HPLC (ThermoScientific Dionex, San Jose, USA) to deliver a flow of approximately 300 nL/min. A C18 Acclaim PepMap100 5 µm, 100 µm x 20 mm nanoViper (ThermoScientific Dionex, San Jose, USA), trapped the peptides prior to separation on a C18 BEH130 1.7 µm, 75 µm x 250 mm analytical UPLC column (Waters, UK). Peptides were eluted with a 60-minute gradient of acetonitrile (2% to 80%). The analytical column outlet was directly interfaced via a nano-flow electrospray ionisation source, with a quadrupole Orbitrap mass spectrometer (Q-Exactive HFX, ThermoScientific). MS data were acquired in data-dependent mode using a top 10 method, where ions with a precursor charge state of 1+ were excluded. High-resolution full scans (R=60,000, m/z 300-1800) were recorded in the Orbitrap followed by higher energy collision dissociation (HCD) (26 % Normalised Collision Energy) of the 10 most intense MS peaks. The fragment ion spectra were acquired at a resolution of 15,000 and dynamic exclusion window of 20s was applied. Raw data files from LC-MS/MS data were processed using Proteome Discoverer v2.1 (Thermo Scientific), and then searched against a human protein database (UniProtKB, reviewed) using the Mascot search engine programme (Matrix Science, UK). Database search parameters were set with a precursor tolerance of 10 ppm and a fragment ion mass tolerance of 0.2 Da. One missed enzyme cleavage was allowed and variable modifications for oxidised methionine, carbamidomethyl cysteine, pyroglutamic acid, phosphorylated serine, threonine and tyrosine were included. MS/MS data were validated using the Scaffold programme (Proteome Software Inc., USA). For the analysis of mass spectral intensities: All raw files were processed with MaxQuant v1.5.5.1 using standard settings and searched against the UniProt Human Reviewed KB with the Andromeda search engine integrated into the MaxQuant software suite (*48*). Enzyme search specificity was Trypsin/P for both endoproteinases. Up to two missed cleavages for each peptide were allowed. Carbamidomethylation of cysteines was set as fixed modification with oxidised methionine and protein N-acetylation considered as variable modifications. The search was performed with an initial mass tolerance of 6 ppm for the precursor ion and 0.5 Da for MS/MS spectra. The false discovery rate was fixed at 1% at the peptide and protein level. Statistical analysis was carried out using the Perseus module of MaxQuant (*49*). Peptides mapped to known contaminants and reverse hits were removed, and only protein groups identified with at least two peptides, one of which was unique, and two quantitation events were considered for data analysis. Each protein had to be detected in at least two out of the three replicates. Missing values were imputed by values simulating noise using the Perseus’ default settings. To calculate p-values, t-tests was performed.

### Expression of ^15^N labelled CPD(1321-1380) for NMR analysis

The GST-CPD(1321-1380) fusion contains a PreScission protease cleavage site downstream of the GST tag. An overnight 2xTY starter culture was inoculated into 1L flasks of NH_4_Cl free M9 medium containing 1.7 g/L of yeast nitrogen base (Sigma Y1251), 1 g/L of ^15^NH_4_Cl (Sigma 299251), and 4 g/L glucose. Protein expression was induced using 300 µM of IPTG and incubated overnight at 16°C for 16 hours, and bacterial lysate prepared as described above for non-isotopically labelled GST fusions. Excess of glutathione-Sepharose beads (500 µL of bead slurry per litre of 2xTY culture) were incubated with the clarified bacterial lysate for 1h at 4°C. Beads were then washed twice with lysis buffer and incubated with the clarified lysate for 1h at 4°C on a rotator. Beads were then washed twice with lysis buffer (50 mM Tris, pH 7.4, 150 mM NaCl, 1 mM EDTA, 1% Triton X-100), and twice with NMR buffer (50 mM Tris, pH 7.4, 150 mM NaCl, 5 mM 2-mercaptoethanol). Beads were then incubated with GST-tagged PreScission protease (GE Healthcare GE27-0843-01) at a concentration of 50 µL of enzyme per 1 mL of bead slurry, and incubated from 5h to overnight at 4°C on a rotator. The supernatant was collected, concentrated using an Amicon 0.5 mL 3,000 NMWL centrifugal filter (Millipore UFC500324), and flash frozen in liquid nitrogen. Protein concentration was measured using Bradford reagent (BioRad 5000006).

### NMR data collection and analysis for CPD

All CPD NMR data sets for were collected at 278K using a 600 MHz Bruker Avance III spectrometer with a 5 mm TCI triple resonance cryoprobe. All samples were prepared with 5% D_2_O as a lock solvent, at pH 7.4, 50 mM Tris, 150 mM NaCl 150 and 5 mM 2-mercaptoethanol. ^1^H-^15^N BEST-TROSY (Band selective Excitation Short Transients-Transverse Relaxation Optimised SpectroscopY) were collected for all samples using an optimised pulse sequence (*50*). The assignment of backbone NH, N, Cα, Cβ and C’ resonances of the 86 μM ^15^N-^13^C CPD (residues 1321-1380) sample was completed using standard 3D datasets acquired as pairs to provide own and preceding carbon connectivities, and between 20-40% non-uniform sampling to aid faster data acquisition. Both the HNCO and HN(CA)CO experimental pair and the CBCA(CO)NH and HNCACB pair were collected with 1024, 64 and 96 complex points in the proton, nitrogen and carbon dimensions respectively. Nitrogen connectivities were established using (H)N(COCA)NNH and (H)N(CA)NNH experiments with 2048, 64 and 96 complex points in the proton, direct and indirect nitrogen dimensions respectively. All data were processed using Topspin versions 3.2 or 4 (Bruker) or, if required, NMRPipe (*51*), with compressed sensing for data reconstruction (*52*), and analysed using NMRFAM-Sparky. The backbone assignment was completed using MARS (*53*). Binding of TBC1D23 to the CPD construct was observed by ^1^H-^15^N BEST-TROSY. Unlabelled TBC1D23 was added to ^15^N labelled CPD with a final concentration of both proteins at 38 μM. The ^1^H-^15^N BEST-TROSY spectrum was compared to a second spectrum recorded for a 38 μM sample of free ^15^N CPD. While some small peak perturbations were observed the main differences caused by TBC1D23 binding were line broadening. This was quantified by taking the peak height ratios of the bound and free spectra. To analyse the chemical shift perturbations the following equation was used to report calculated the distance between each peak in the bound spectrum when compared to the assigned, free CPD: Δδ“total”= ((δH)^2+ (δN/5)^2)^0.5, With the smallest value reported as the minimal chemical shift perturbation.

### Isothermal Titration Microcalorimetry

Experiments were performed using a Nano ITC machine from TA Instruments. TBC1D23 C-terminal domains were gel filtered into buffer composed of 100 mM NaCl, 100 mM Tris pH 7.4 and 1 mM TCEP. Peptides were dissolved in the same batch of buffer. Both the WT and mutant C-terminal domains were concentrated to 100 μM (1.4 mg/ml) with peptide concentration varying between 0.8 and 5 mM (depending on a peptide). Peptides were titrated into TBC1D23 C-terminal domains with 20 injections of 2.4 μl each separated by 300 second intervals. Experiments were conducted at 20°C. A relevant syringe-peptide-into-buffer blank was subtracted from all data and for constructs which displayed measurable binding, three independent runs that showed clear saturation of binding were used to calculate the mean K_D_ of the reaction, the stoichiometry (n), and the corresponding SEM values using NanoAnalyze. Both curves and trace data were subsequently exported for figure generation in Prism.

### Protein expression and purification for structural analysis

Recombinant proteins were expressed in BL21 plyS *E. coli* grown in shaking 2xTY media at 37°C. Expression was induced with 0.2 mM IPTG and cells grown overnight at 22°C. Cells were resuspended in buffer A (250 mM NaCl, 20 mM Tris (pH 7.4), 1 mM DTT) supplemented with AEBSF hydrochloride, MnCl_2_ and DNAse1. Cell pellets were lysed using a cell disruptor (Constant Systems) before clarification by ultracentrifugation (104350 RCF) for 45 minutes. The supernatant was batch bound to glutathione Sepharose resin (Cytiva) in buffer A before washing with 400 ml of buffer A and cleavage of the GST-tag with overnight at room temperature with thrombin. The resultant flow through was concentrated for gel filtration Superdex 200 column (GE Healthcare) into either buffer A for crystallization or into ITC buffer (100 mM NaCl, 100 mM Tris (pH 7.4) and 1 mM TCEP).

### X-ray crystallography

TBC1D23 C-terminal domains was concentrated to either 11 mg/ml (full length) or 14 mg/ml (-VLDALES) and mixed with 1.5 times molar excess syntaxin-16_209-221_ peptide. High throughput sitting drops were used to obtain crystallisation conditions, which were further optimisation before being repeated in hanging drop with protein:peptide complex being mixed 1:1 ratio with crystallisation mother liquor. TBC1D23 full-length C-terminal domain was crystallised in 0.02 M citric acid, 0.08 M Bis-Tris propane pH 8.8 and 16% w/v PEG3350. TBC1D23 (-VLDALES) C-terminal domain was crystallised in 0.8 M potassium phosphate dibasic, 0.1 M HEPES/NaOH pH 7.5, 0.8 M sodium phosphate monobasic and 1% 1,2-butandiol. Crystals were cryoprotected by soaking in mother liquor supplemented with 35% glycerol and 1 mg/ml syntaxin-16_209-221_ peptide and flash cooled in liquid nitrogen.

Diffraction data were collected at Diamond Light Source on the IO4 beamline at 100K and were processed with Autoproc. Structures were solved by molecular replacement with PHASER MR using a single copy of the TBC1D23 C-terminal domain of (6JM5) with zinc and water molecules removed as a search model. REFMAC5 was used for iterative rounds of refinement interspersed by manual rebuilding of the model using COOT. Crystallographic programs were ran using the CCP4i2 package and figures were rendered using UCSF ChimeraX (*54*). Data collection and refinement statistics are summarised in Table S1. Electron density of the full length TBC1D23 domain structure in the presence of syntaxin-16 peptide clearly showed C terminal tails bound in two molecules, syntaxin-16 bound in a third with the fourth being somewhat ambiguous (fig. S4). Deletion of the C-terminal tail (VLDALES) and inclusion of syntaxin-16 peptide resulted in a structure whose electron density when solved by MR using 6JM5 minus the VLDALES as the search model, unambiguously showed the syntaxin-16 peptide bound in all molecules.

Coordinates and MTZ files are deposited on PDB under accession codes (PDB:XXXX PDB:XXXX). replication of the results. Begin with a section titled Experimental Design describing the objectives and design of the study as well as prespecified components.

In addition, include a section titled Statistical Analysis at the end that fully describes the statistical methods with enough detail to enable a knowledgeable reader with access to the original data to verify the results. The values for *N*, *P*, and the specific statistical test performed for each experiment should be included in the appropriate figure legend or main text.

All descriptions of materials and methods should be included after the Discussion. This section should be broken up by short subheadings. Under exceptional circumstances, when a particularly lengthy description is required, a portion of the Materials and Methods can be included in the Supplementary Materials.

## Acknowledgments

We thank Michael Ailion, Dan Billadeau, Lloyd Fricker, and Olga Perisic for providing reagents, Jérôme Boulanger for help with image analysis, Sally Gray for technical assistance, and Airlie McCoy for invaluable help compiling the structural data. We thank Diamond Light Source for beamtime and the staff of beamline IO4 for assistance with crystal testing and data collection.

## Funding

Medical Research Council, as part of United Kingdom Research and Innovation (also known as UK Research and Innovation) file reference number MC_U105178783 (SM) Wellcome Trust grant number 207455/Z/17/Z (DJO)

## Author contributions

Conceptualization: SM

Methodology: TJS

Investigation: JCO, JGGK, AKG, JLW, SYPC, DJO

Supervision: SM, DJO

Writing—original draft: JCO, SM, DJO, JLW

Writing—review & editing: SM, DJO

## Competing interests

Authors declare that they have no competing interests.

## Data and materials availability

All data are available in the main text or the supplementary materials.

## Supplementary Materials

**This PDF file includes:**

Tables S1 and S2

Figs. S1 to S5

**Other Supplementary Material for this manuscript includes the following:**

Data S1. Mass spectrometry data as plotted in Figs. 1, B and C; and Fig. 3D.

Data S2. Source data for plots shown in Fig. 2, B, E and F; Fig. 3I; fig. S1, C to E; and fig S3G.

Data S3. Antibodies, plasmids and cell lines used in this study.

## Supplementary Figure Legends

**Fig. S1.**
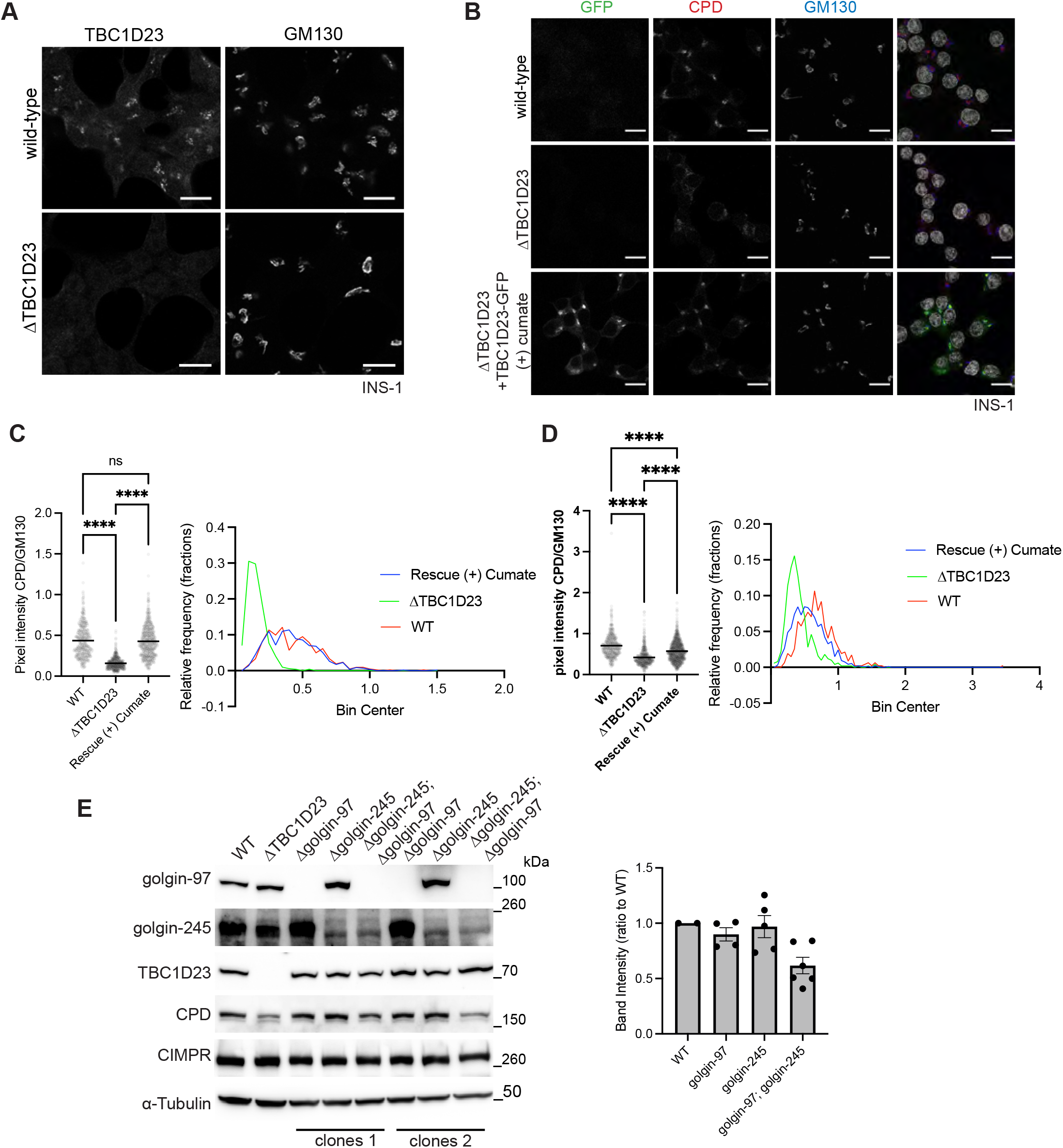
Normal CPD trafficking requires TBC1D23. (A) Confocal micrographs of wild-type and ΔTBC1D23 INS-1 cells, labelled for endogenous TBC1D23 and GM130 (Golgi). Scale bar: 10 µm. (B) Confocal micrographs of the indicated INS-1 cell lines, labelled for GFP-booster and endogenous CPD and GM130 (Golgi). (C) Scatter plot showing the ratio of the Golgi-area fluorescence intensity of CPD over GM130 for cells as in (B). Golgi-area were detected automatically using an ImageJ macro (see Materials and Methods). The black bar is mean, and for wild type (WT), n = 422 cells, for ΔTBC1D23, n=521, rescue, n=573. Statistics: ordinary one-way ANOVA followed by a Sidak’s multiple comparison tests with a single pooled variance. ****: P<0.0001, ns: not significant. Also shown are the ratios plotted as frequency distributions with a bin width of 0.05 (see material and methods). (D) An independent biological repeat of the experiment shown in (c). For wild type (WT), n = 468, for ΔTBC1D23, n = 501, for the rescue, n = 946. (E) Immunoblots of detergent lysates from HEK-293 WT, ΔTBC1D23, 2 clones of Δgolgin-97, two clones of Δgolgin-245 and two clones of Δgolgin-97;Δgolgin-245. Total protein concentration was measured so equal amount of protein was loaded in each lane with band intensity quantification of the CPD blot normalised to WT for each blot. Source data for fig. S1, C to E are in data S2.

**Fig. S2.**
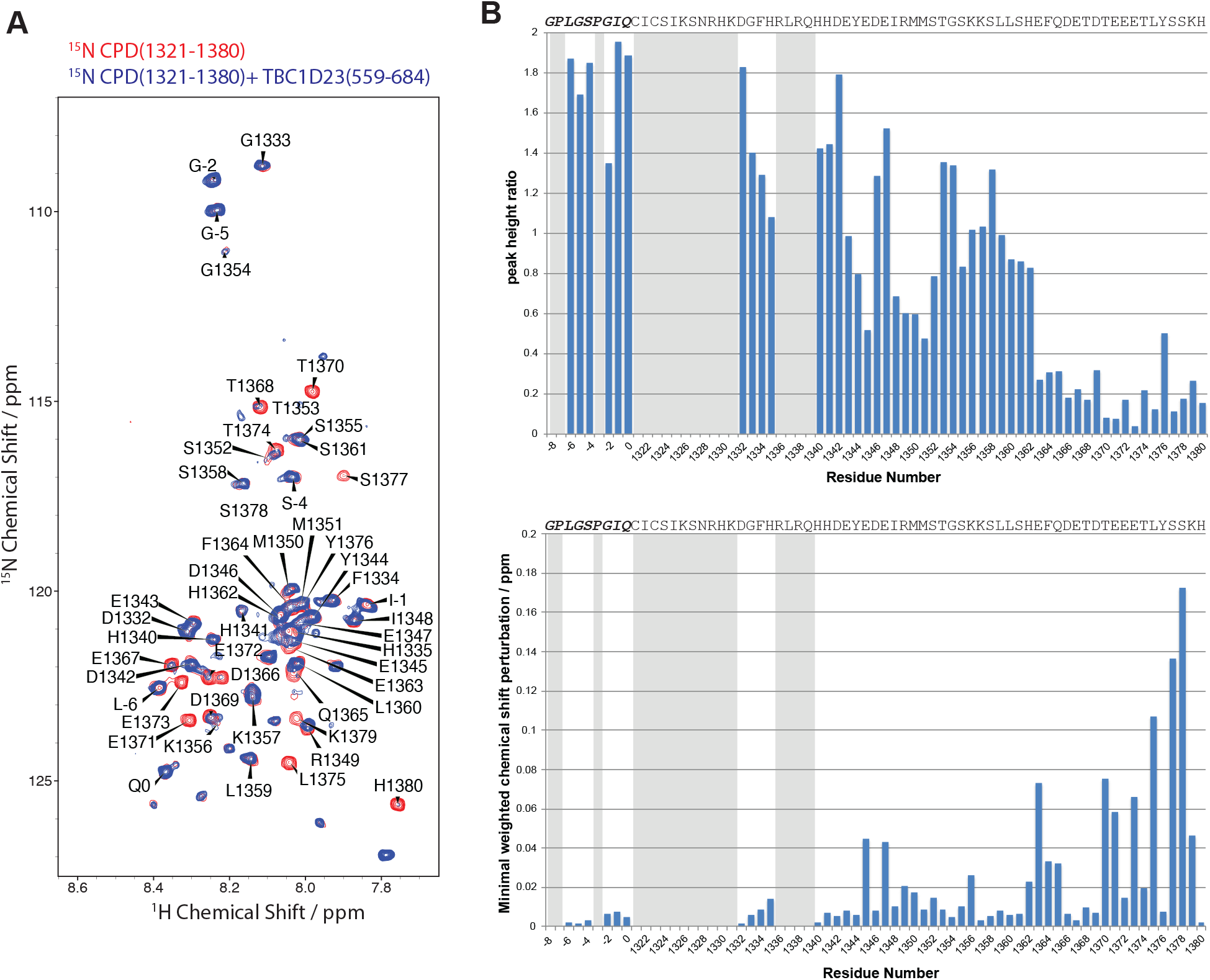
NMR analysis of CPD cytoplasmic tail in presence of TBC1D23 C-terminal domain. (A) The assigned ^1^H-^15^N BEST-TROSY of ^15^N labelled CPD (residues 1321-1380) is shown in red overlaid with the BEST-TROSY of ^15^N CPD with unlabelled TBC1D23 (residues 559-684) at a 1:1 ratio shown in blue. Both spectra were collected at 278K and 600 MHz. (B) Peak height intensity differences of the ^15^N CPD peaks in the presence and absence of unlabelled TBC1D23 are reported as a ratio (top panel). The minimal chemical shift perturbations seen for ^15^N CPD in the presence and absence of unlabelled TBC1D23 (bottom panel). Greyed areas indicate missing assignments.

**Fig. S3.**
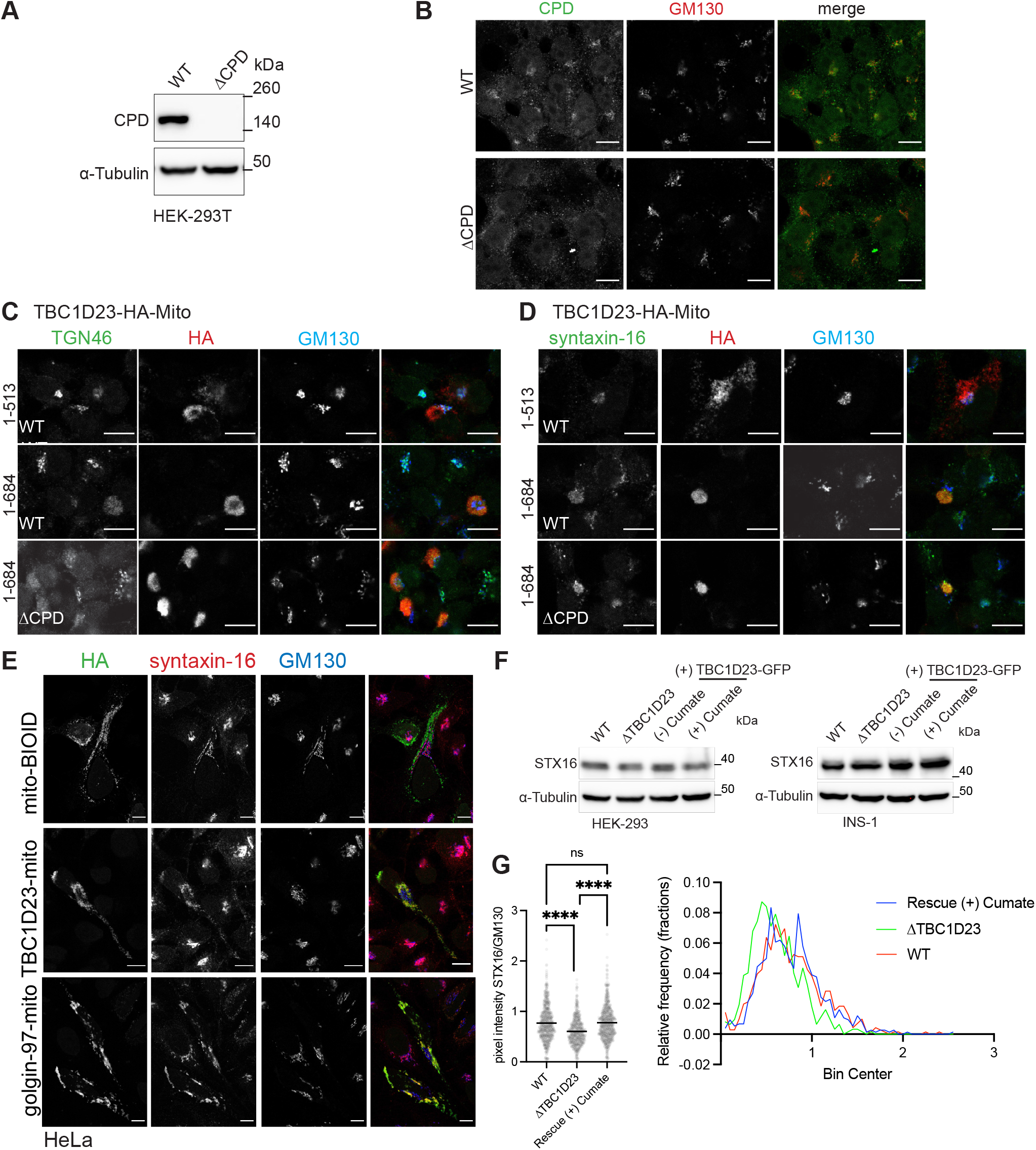
CPD is not required for TBC1D23 to capture vesicles. (A) Immunoblots of detergent lysates from HEK-293T WT and ΔCPD. Cells were lysed with Triton-X-100. Total protein concentration was measured so equal amount of protein was loaded in each lane. (bottom) band intensity quantification of the CPD blot normalised to WT for each blot. (B) Confocal micrographs of WT and ΔCPD HEK-293T cells. CPD was knocked out using CRISPR/Cas9. Cells were fixed, permeabilised and stained using endogenous CPD and GM130 (Golgi marker). Scale bar: 10 µm. (C) Confocal micrographs of HeLa cells TBC1D23-mito chimaeras and labelled for the HA tag in the chimaera, endogenous TGN46 and GM130. Scale bar: 10 µm. (D) Confocal micrographs of HeLa cells TBC1D23-mito chimaeras and labelled for the HA tag in the chimaera, endogenous syntaxin-16 and GM130. Scale bar: 10 µm. (E) Confocal micrographs of HeLa cells expressing golgin-97-mito or TBC1D23-mito chimaeras and labelled for the HA tag in the chimaera and endogenous syntaxin-16 and GM130. Mito-BioID was used as a negative control. Scale bar: 10 µm. (F) Immunoblots of detergent lysates from HEK-293 or INS-1 cells as indicated. In each case WT, ΔTBC1D23 and rescue ΔTBC1D23 cells stably expressed TBC1D23-GFP under a cumate promoter, and the cells were incubated in the absence (-) and in presence (+) of cumate for 24-36 hours and lysed with Triton-X-100. Total protein concentration was measured so equal amount of protein was loaded in each lane. These are the samples as in Fig. 2A, but here are blotted for syntaxin-16. (G) As Fig. 3I: independent biological repeat for syntaxin-16 Golgi localisation. For WT, n = 705, for ΔTBC1D23, n = 619, for the stable rescues, n = 743. Source data for fig. S3G are in data S2.

**Fig. S4.**
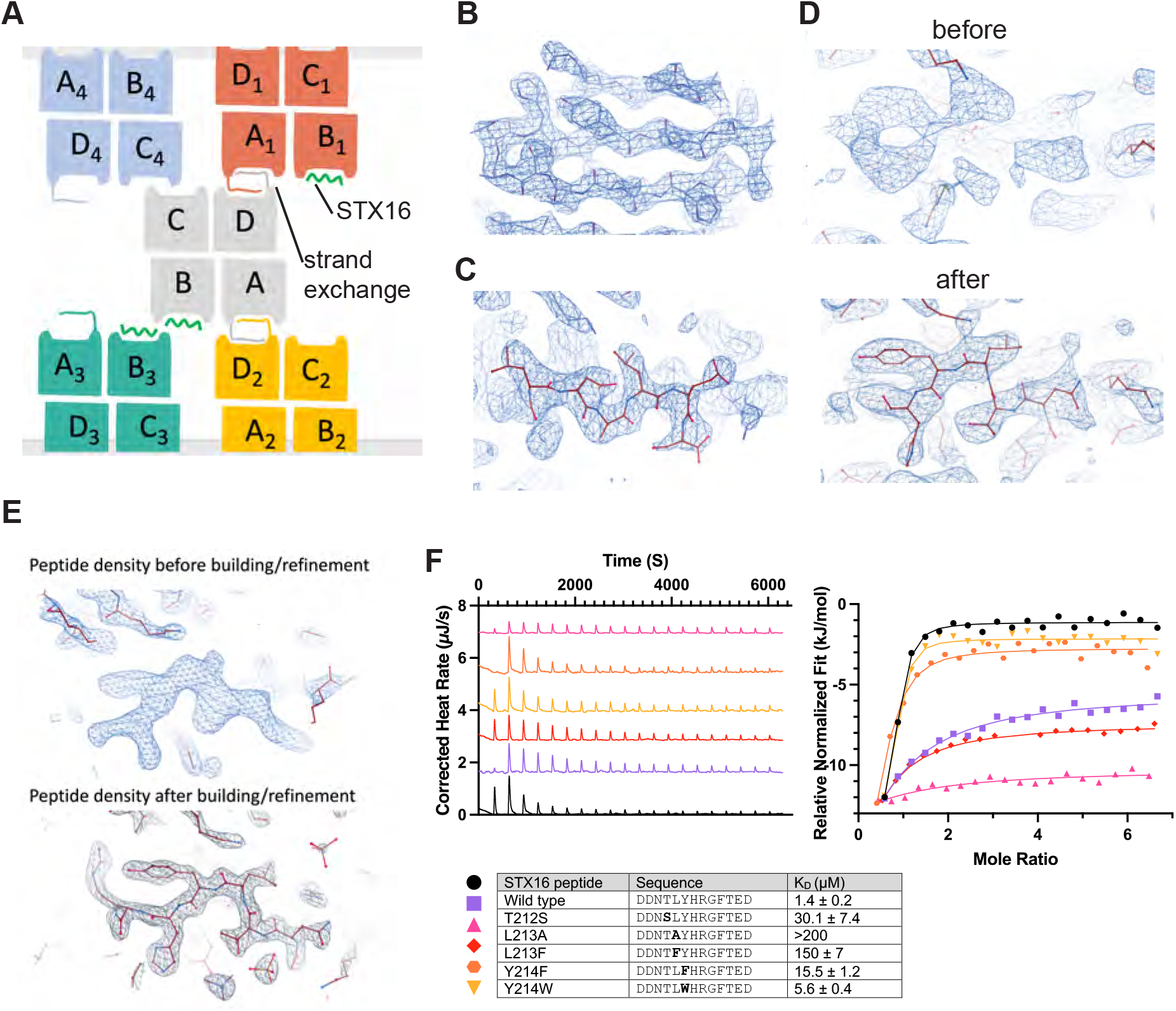
Structural and biochemical characterisation of the interaction between the TBC1D23 C-terminal domain and the syntaxin-16 peptide. (A) Cartoon of the crystal packing of the four copies of the TBC1D23 C-terminal domain showing the C-terminal strand exchange that occurs for two of the copies A and D, with the syntaxin-16 peptide bound to copy B, and the occupancy of copy C unclear. (B) Representative electron density of full-length TBC1D23 C-terminal domain contoured at 2σ. (C) Electron density of TBC1D23 full length tail (..MKVLDALES) blocking the syntaxin-16 binding site, as seen for copy C and D in (A). (D) Electron density of the syntaxin-16 peptide bound to the TBC1D23 full length C-terminal domain (copy B in (A)), shown before and after building and refinement. (E) Density map of showing the initial experimental density (blue) obtained with MR for the C-terminal lacking the last eight residues domain with syntaxin-16 peptide bound, and the final 2Fo-Fc electron density (black) with syntaxin-16 peptide built in. (F) ITC of relative binding of mutated forms of the syntaxin-16 peptide to TBC1D23 C-terminal domain. showing range of similar residues can facilitate binding.

**Fig. S5.**
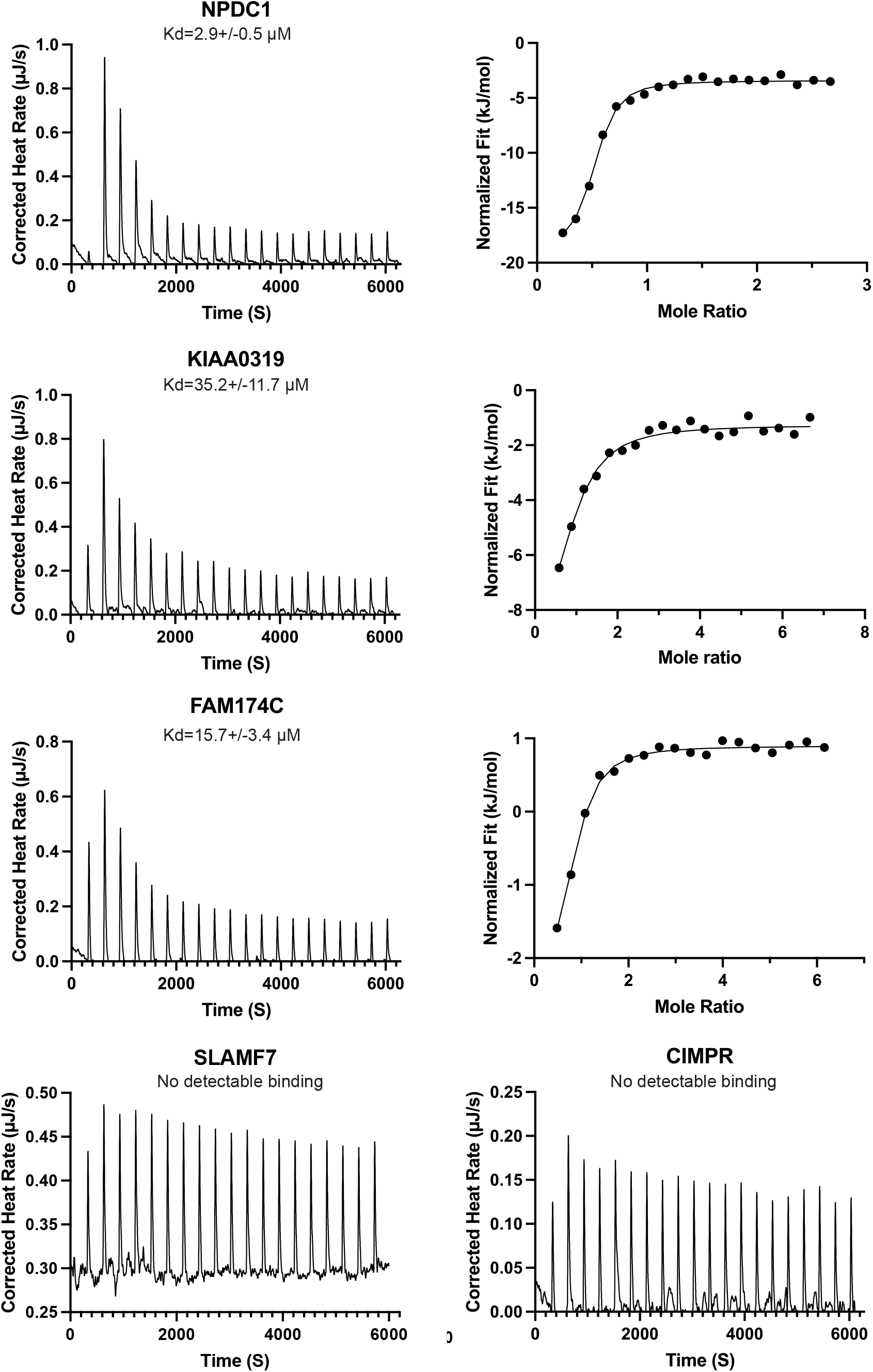
Binding of peptides containing the acidic TLY motif from several proteins that recycle between endosomes and Golgi. ITC of the binding to the TBC1D23 C-terminal domain of peptides corresponding to TLY-motif containing regions from the cytoplasmic tails of the indicated proteins.

**Table S1.** Crystallographic data collection and refinement statistics.

|  | **PDB:XXXX1** | **PDB:XXX2** |
| --- | --- | --- |
| Space group | **P 62 2 2** | **P 65 2 2** |
| Resolution range | 194.38-3.17 | 116.19-2.19 |
| InnerShell | 112.394-11.533 | 116.19-8.280 |
| OuterShell | 3.814-3.172 | 2.479-2.187 |
| Rmerge (all I+ & I-) | 0.939 (2.988) | 0.133 (2.607) |
| Rmerge (within I+ & I-) | 0.989 (2.998) | 0.141 (2.710) |
| Rmeas (all I+ & I-) | 0.951 (3.028) | 0.136 (2.651) |
| Rmeas (within I+ & I-) | 1.013 (3.073) | 0.145 (2.791) |
| Rpim (all I+ & I-) | 0.151 (0.485) | 0.024 (0.472) |
| Rpim (within I+ & I-) | 0.220 (0.667) | 0.034 (0.668) |
| Number of observations | 352792 (17313) | 486437 (23072) |
| Number of unique observations | 9118 (457) | 14895 (745) |
| <I>/sd(I)> | 7.5 (1.9) | 23.0 (1.9) |
| CC(1/2) | 0.972 (0.627) | 0.999 (0.707) |
| Completeness % (spherical) | 90.8 (47.7) | 43.2 (7.0) |
| Completeness % (ellipsoidal) | 90.8 (47.7) | 94.8 (82.9) |
| Multiplicity | 38.7 (37.9) | 32.7 (31.0) |
| RMS bond lengths (Å) | 0.0063 | 0.0077 |
| RMS bond angles (°) | 1.644 | 1.605 |
| Reflections (All/Free) | 8038 / 356 | 14895/752 |
| R<sub>factor</sub>/R<sub>free</sub> | 0.209 / 0.274 | 0.186 / 0.259 |
| Ramachandran (%) (favoured/outliers) | 97.11/2.89 | 99.24/0.76 |

**Table S2.**
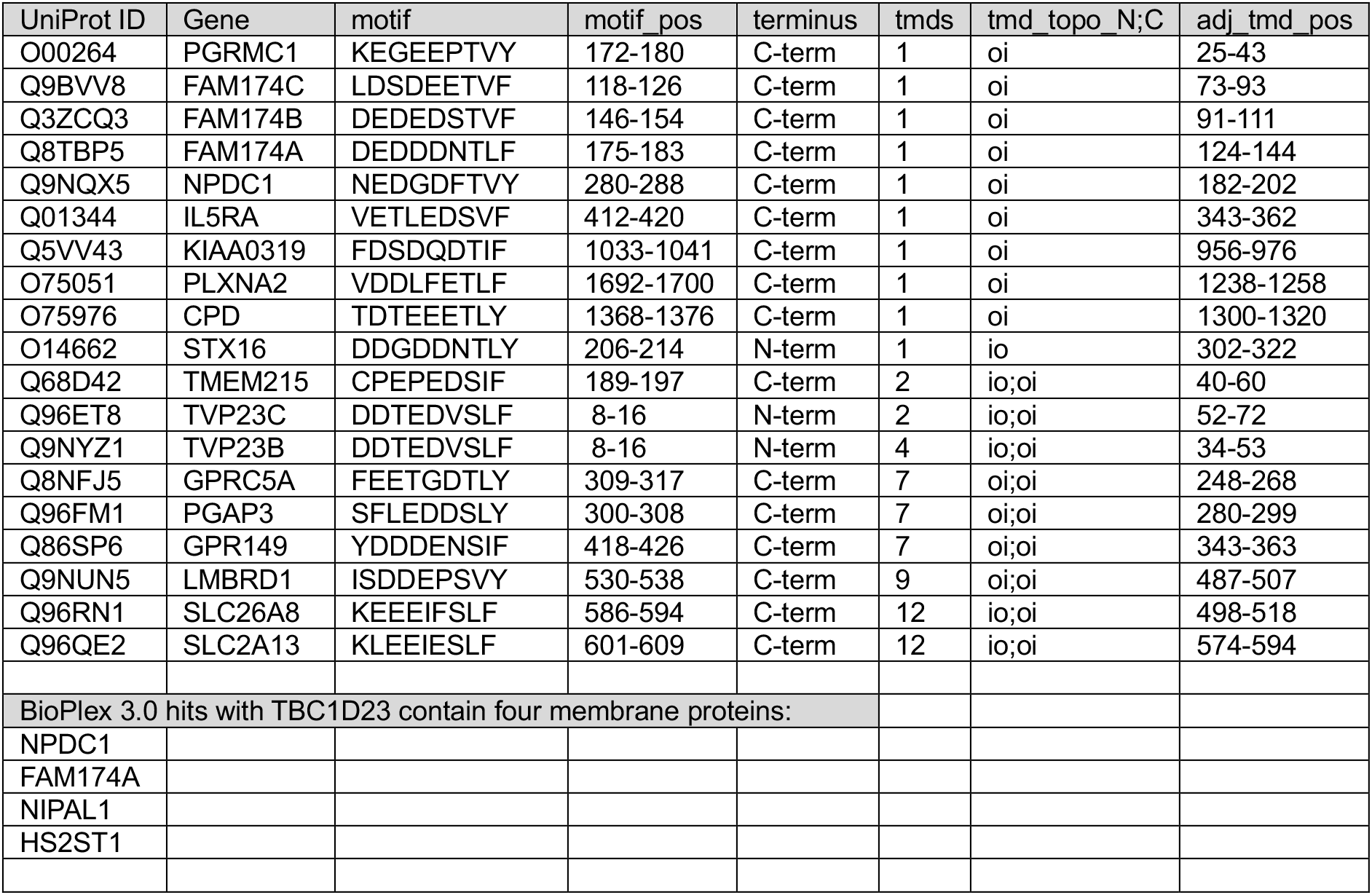
Human membrane proteins with the motif [DE]_>=3/5_[TS][LIV][YFW] in a cytoplasmic domain. Also shown are the four membrane proteins reported to interact with TBC1D23 in the BioPlex protein interaction screen.

## Notes

### Competing Interest Statement

The authors have declared no competing interest.

